# Seq2Karyotype (S2K): A Method for *in-silico* Karyotyping Using Single-Sample Whole-Genome Sequencing Data

**DOI:** 10.1101/2025.08.25.671339

**Authors:** Limeng Pu, Karol Szlachta, Virginia Valentine, Meiling Jin, Sivaraman Natarajan, Robert Greenhalgh, Nadezhda V. Terekhanova, Jian Wang, Daniel Putnam, Li Dong, Lingxiang Jiang, Soumya Tumbath, Xiumei Huang, Xiaotu Ma, Thomas Look, Marcin Wlodarski, Lu Wang, Steven Burden, John Easton, Xiang Chen, Jinghui Zhang

## Abstract

DNA abnormalities characterized by cytogenetic imaging at the single cell resolution, i.e. karyotyping, have long served as cancer diagnostic and prognostic biomarkers. To enable *in-silico* karyotyping using unpaired whole-genome sequencing data, we developed Seq2Karyotype (S2K), a tool that fits karyotype models with clonality estimation based on read-depth and allelic imbalance in a bulk sample and supports visualization-guided refinement. Analysis on 19 adult and pediatric cancer cell lines revealed unexpected intratumoral heterogeneity involving multiple copy number variation (CNV) states including whole-genome duplication, which were validated by imaging and single-cell omics profiling. Analyses on patient samples showed high concordance with clinical cytogenetic reports for acute myeloid leukemia, and revealed evolutionary trajectories from multi-region metastatic neuroblastomas implicating reversion. These findings highlight extensive and dynamic intratumoral heterogeneity contributed by CNV in both cell line models and patient samples, which may inform future research on tumor evolution under selective pressure such as drug exposure.

## Introduction

Analysis of copy number variations (CNVs) in tumor and germline samples are important for discovery of predisposition events [1, 2] and risk stratification for precision treatment in cancer and other diseases [3, 4]. Cytogenetic techniques such as Fluorescence In Situ Hybridization (FISH) and Spectral Karyotyping (SKY) mapping play a vital role in this process. These techniques are used to examine individual cells, thus the resulting data also provide insights on intratumoral heterogeneity (ITH) and clonal evolution [5]. As next-generation sequencing (NGS) has become routine for both research and clinical testing, methods have been developed to integrate read depth-based changes and allelic imbalance to determine the CNV status and to impute tumor heterogeneity. For example, ASCAT [6] and ABSOLUTE [7] are widely used for estimating purity, ploidy, and absolute copy numbers in tumor samples. TITAN [8] extends this by modeling ITH and clonal expansions.

Existing methods implementing allelic imbalance analysis have several major limitations. First, most of these tools require paired tumor-normal data, where heterozygous single nucleotide polymorphisms (SNPs) in the matched normal sample are used to evaluate allelic imbalance in the matched tumor, helping mitigate bias and artifacts in the analysis [9]. However, paired tumor-normal CNV analysis is infeasible under several important circumstances: 1) established cancer cell lines are used extensively as model systems for research and drug screening but most of them lack paired normal samples; 2) germline samples may contain somatically acquired CNVs that contribute to cancer initiation and progression, such as mosaic monosomy 7 or uniparental disomy (UPD) in hematopoietic malignancies [10]; and 3) tumor-only CNV analysis may be performed for time-critical clinical decision-making, as access to germline samples often requires additional time to acquire consent and prepare samples [11]. Another major limitation is the lack of modeling of multiple CNV states as existing methods only consider subclonal presence of single CNV events. For example, single-cell whole-genome sequencing (scWGS) of COLO829, a melanoma cell line commonly used as a standard for somatic variant detection, showed co-existence of cells with monosomic and trisomic karyotypes which were rarely captured by existing CNV analysis tools [12]. Our prior study also revealed co-existence of subclones harboring different CNV states of the same region in multi-region autopsy samples of diffuse midline gliomas treated with a *PDGFR* inhibitor [13].

To overcome these limitations, we developed Seq2Karyotype (S2K), a method that uses whole-genome sequencing (WGS) data generated from a single, unpaired sample to model CNV heterogeneity, considering the possibility of multiple CNV events at the same genomic locus (**Fig. 1A**). We demonstrate S2K’s capability in modeling ITH by analyzing four bulk WGS datasets generated from adult cancer cell lines commonly used for benchmark analyses, pediatric neuroblastoma cell lines, patient acute myeloid leukemia (AML) samples and multi-region metastatic neuroblastoma (NBL) samples (**Fig. 1B**). The subclonal and/or complex admixed CNV events detected by S2K, which include whole-genome duplications (WGDs), unveiled extensive and evolving ITH in both cell lines and patient tumors. These results, validated by karyotyping, FISH, SKY mapping, scWGS and scRNA-seq (**Fig. 1B**), highlight the power of S2K in unveiling the significant contribution of CNV to ITH which may inform future studies on tumor evolution.

**Figure 1.**
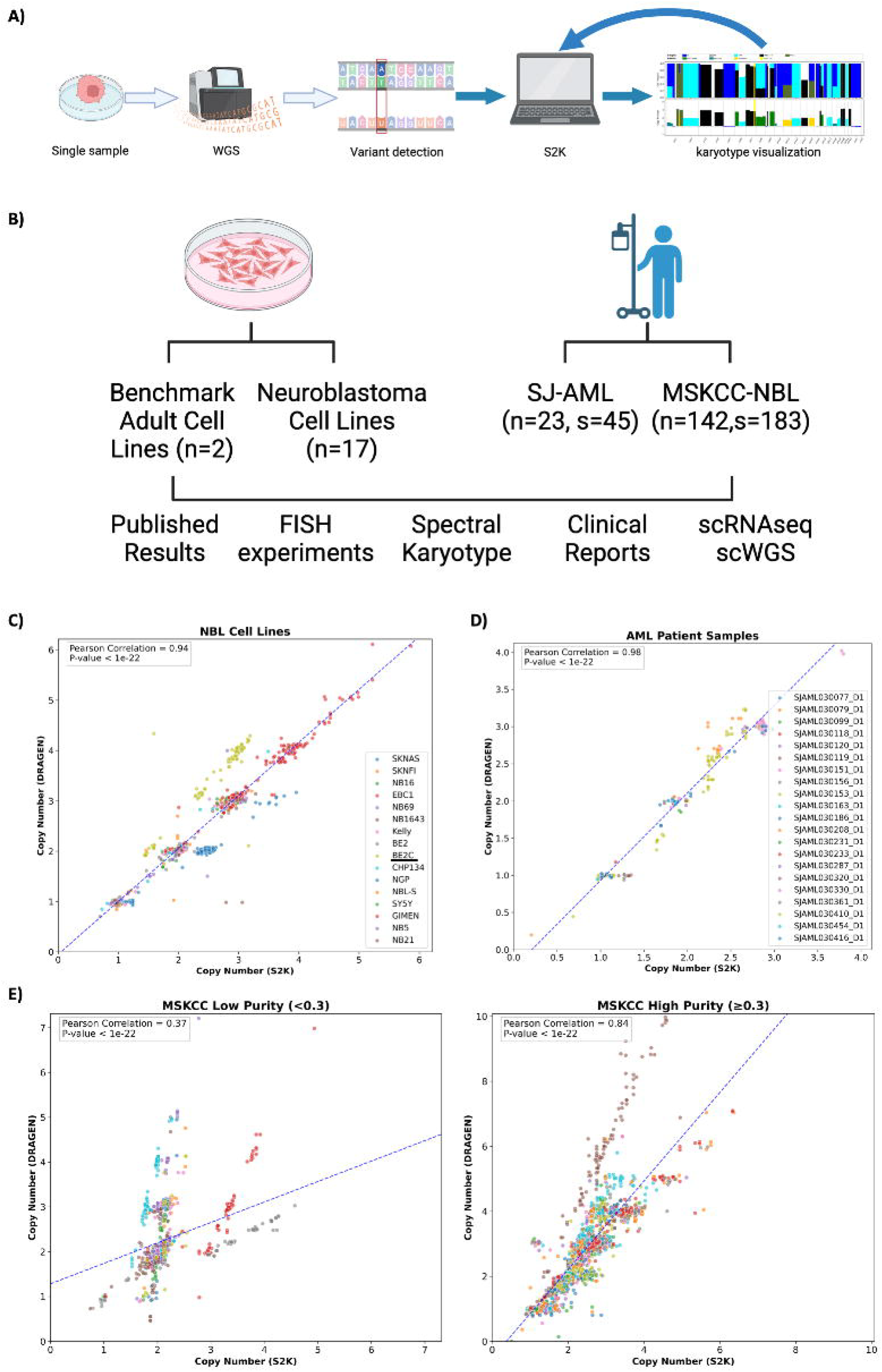

## Results

### 1. S2K workflow

S2K streamlines six data analysis steps followed by visualization of predicted karyotypes and their clonality (**Supplementary Fig. S1A)**. To enable single-sample analysis, S2K identifies putative heterozygous SNPs in a single genome from a high-quality SNP list constructed from ∼8,000 germline WGS samples (**Supplementary Information**). Designed for detecting potentially subclonal gross CNVs, S2K focuses on events spanning over 1Mb, finding the best-matched model with potentially mixed populations of CNVs and their associated clonality, similar to a karyotype report (**Supplementary Fig. S1B)**. The results can be visualized for further model refinement, with an option to recalibrate the reference diploid region. Details are described in **Methods, Supplementary Information**, and **Supplementary Fig. S1**.

### 2. CNV state and segmentation performance comparison

As CNV segmentation and state are critical for subsequent karyotype modeling, we compared the corresponding results of S2K with DRAGEN, a tool ranked as the top performer in a recent benchmark analysis using data generated from a series of dilution experiments [14] on the four WGS data sets tested in this study. This comparison does not involve the analysis of CNV heterogeneity, as DRAGEN does not report multiple CNV states at the same locus. In regions where S2K identified multiple CNV states, the predominant CNV state was used for comparison with DRAGEN. The criteria for segmentation comparison and an example of assigning match vs. mismatch in regions where S2K called admixed karyotypes is shown in **Supplementary Fig. S2** using data generated from COLO829 cell line. The overall matching rate for the two benchmark cancer cell lines, COLO829 and HCC1395, and their matched normal, is 91-96%.

For the 17 neuroblastoma (NBL) cell lines analyzed in this study, the copy number data computed by the two methods were highly correlated (R=0.94 **Fig. 1C**), except for BE2C (**Fig. 1C**, yellow dots), which S2K projected to have admixed hypodiploid and hypotetraploid subpopulations, while DRAGEN projected a single tetraploid population. The overall segment matching rate of the two methods is 91% for this cohort (**Supplementary Fig. S3A**). Patient AML samples had a near perfect correlation of 0.98 between CNV data analyzed by S2K and DRAGEN (**Fig. 1D**) and a 98% match on CNV segments (**Supplementary Fig. S3B**). We separated NBL patient samples into low- and high-purity groups using the published tumor purity cutoff of 0.3, based on clinical practice [15]. The overall correlation of high-purity samples (0.84) is significantly higher than that of low-purity samples (0.37) (p < 10^-6^, Wilcoxon’s rank sum test, **Fig. 1E**). The overall CNV segment matching rate is 80% (**Supplementary Fig. S3C**) with significant deviations found in three low-purity samples and one high-purity sample due to discrepancies in diploid reference selection. Tumor purity estimated by DRAGEN and S2K showed high concordance with the published tumor purity of this cohort (**Supplementary Fig. S3D**). However, DRAGEN failed in purity assessment on 12 samples (Supplementary Fig. S**3D, left)** which had lower purity compared to the remaining samples (mean: 0.21 vs. 0.56, p = 0.0028, Wilcoxon’s rank sum test).

### 3. Evolving clonal heterogeneity in benchmark cell lines

The karyotype modeling implemented in S2K can unveil ITH resulting from an admixed CNV state. We first evaluated the S2K results on COLO829 (melanoma) and HCC1395 (breast cancer), two adult cancer cell lines commonly used as a community reference for characterizing complex and aneuploid genomes [16, 17] owing to the availability of their matched normal cells for somatic variant detection in paired tumor-normal analysis. The public availability of scWGS data as well as karyotyping by SKY mapping [12, 17] can serve as ground truth for assessing accuracy.

In COLO829, S2K analysis revealed complex karyotypes with triploid, tetraploid and diploid copy-neutral LOH (cn-LOH) chromosomes (**Fig. 2A**), consistent with previously published findings by SKY mapping and scWGS [12] albeit with different cancer cellular fraction (CCF) (**Supplementary Fig. S4**). For example, three regions (1p, 10p and chr18) contained an admixture of cn-LOH and 3-copy karyotypes at an approximately 2:1 ratio, with interconnected structural variation (SV) breakpoints (shown as triangles in the top panel of **Fig. 2A**, details in **Supplementary Table 1**). The 3-copy cells matched the profile expected from a previously reported complex derivative chromosome (der18, **Fig. 2B**) [12] which was lost in the subset of cells exhibiting cn-LOH. The genome-wide karyotype is consistent with the evolution model constructed from scWGS [12] which showed that derivative chromosomes (der18 and der7, **Fig. 2B, C**) were formed prior to (partial) WGD, leading to tetraploidy or cn-LOH across the genome. The subsequent loss of der18, consistent with the pattern of reverse evolution, forms a lineage that was the predominant population with a CCF of 0.67 in the COLO829 sample analyzed in this study (**Fig. 2C**) which had varying CCF (e.g. 0.11 in prior scWGS analysis) in samples used in other published studies (**Supplementary Fig. S4, [12]**). No abnormal karyotype was detected for its matching normal sample COLO829BL.

**Figure 2.**
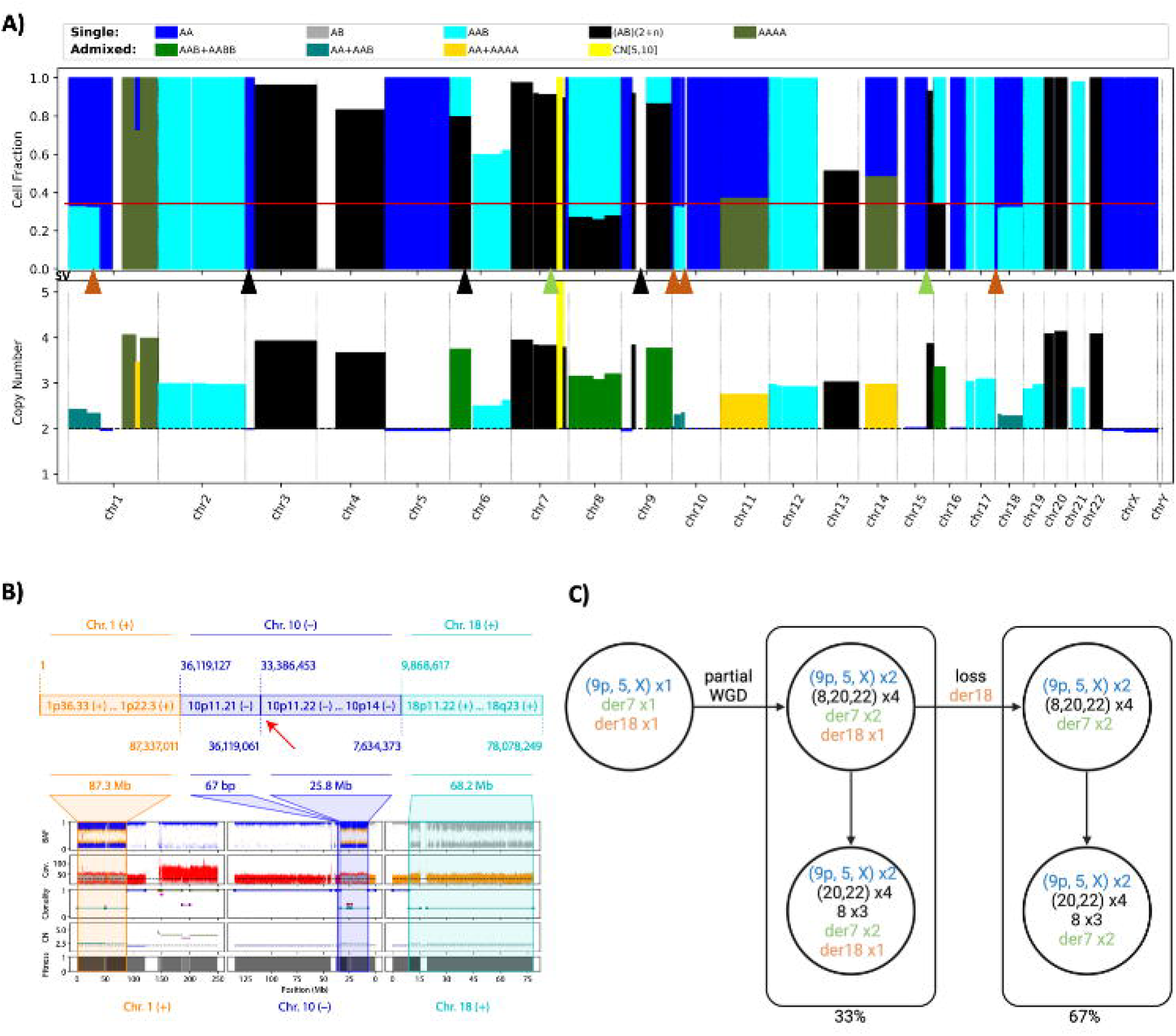

The analysis of the HCC1395 cancer cell line showed admixed karyotypes in multiple chromosomes, consistent with previously published findings by scWGS [17], and the diploid regions are mostly cn-LOH events (**Supplementary Fig. S5**). Unexpectedly, the analysis of its matching normal, HCC1395BL, revealed six gross CNV events including three known clonal losses of 6p, 16q, and X [17] as well as three previously unreported CNV events involving loss of 11q and 17p, and gain of chr20, all with an estimated CCF of 0.7 (**Fig. 3A**). These findings indicate that this normal cell line profiled in our study was composed of two subpopulations: a major subclone containing all six CNVs, and a minor subclone containing only the three known CNVs. To validate this unexpected finding, we performed two cytogenetic experiments on HCC195BL (**Fig. 3B, C**): 1) FISH experiments validated 1-copy gain of chr20 in 81% of the 500 interphase nuclei scored; and 2) SKY mapping confirmed the existence of a major subclone (83% of the cells) harboring all six CNVs and a minor subclone (17%) with only the clonal loss of 6p, 16q and X. SKY mapping also showed that all subarm losses were caused by unbalanced inter-chromosomal translocations with complex re-arrangements from WGS SV analysis (**Fig. 3D** and **Supplementary Table S1**).

**Figure 3.**
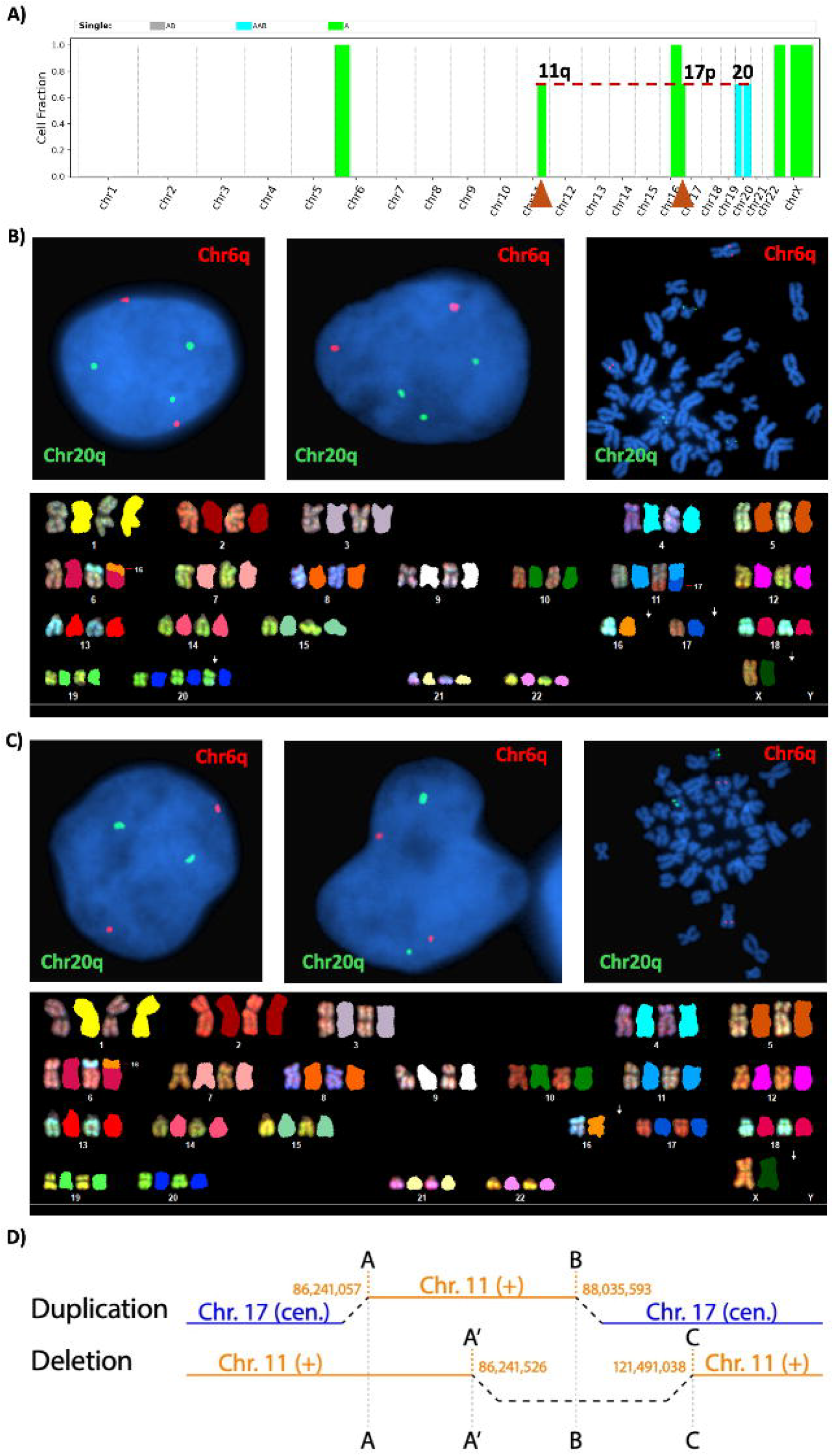

### 4. Intratumoral heterogeneity and evolution in pediatric neuroblastoma cell lines

Unlike adult community reference cell lines (e.g. COLO829 and HCC1395), pediatric cancer cell lines often lack matched normal samples. We analyzed 17 neuroblastoma cell lines, as neuroblastomas are known for complex genomic structures [18]. Overall, S2K’s karyotype modeling projected multiple subclones in 16 out of the 17 lines. We selected SY5Y, BE2, BE2C and NB5 (Supplementary Information) for validation as they contain challenging events such as complex karyotypes or subclones with very low CCF (**Supplementary Tables 2, 3**).

In SY5Y, a cn-LOH event at 4p with an estimated CCF of 0.10 and a 1-copy gain of 17q with an estimated CCF of 0.90 were detected (**Fig. 4A**). Thus, rare subclones were expected due to the presence of a low-CCF event or the absence of a high-CCF event. Indeed, the 17q gain was detected in 91% of cells by FISH (**Fig. 4B**). Since cn-LOH events cannot be validated cytogenetically, we performed scWGS at ∼10X coverage (Methods, **Fig. 4C**). Amongst the 248 cells passing QC checks, 85% exhibited allelic imbalance with diploidy at 4p (**Fig. 4D)**, validating the presence of a subclone with 4p cn-LOH. The high CCF in scWGS analysis indicates a possible selection bias during the scWGS sample preparation (**Fig. 4C**).

**Figure 4.**
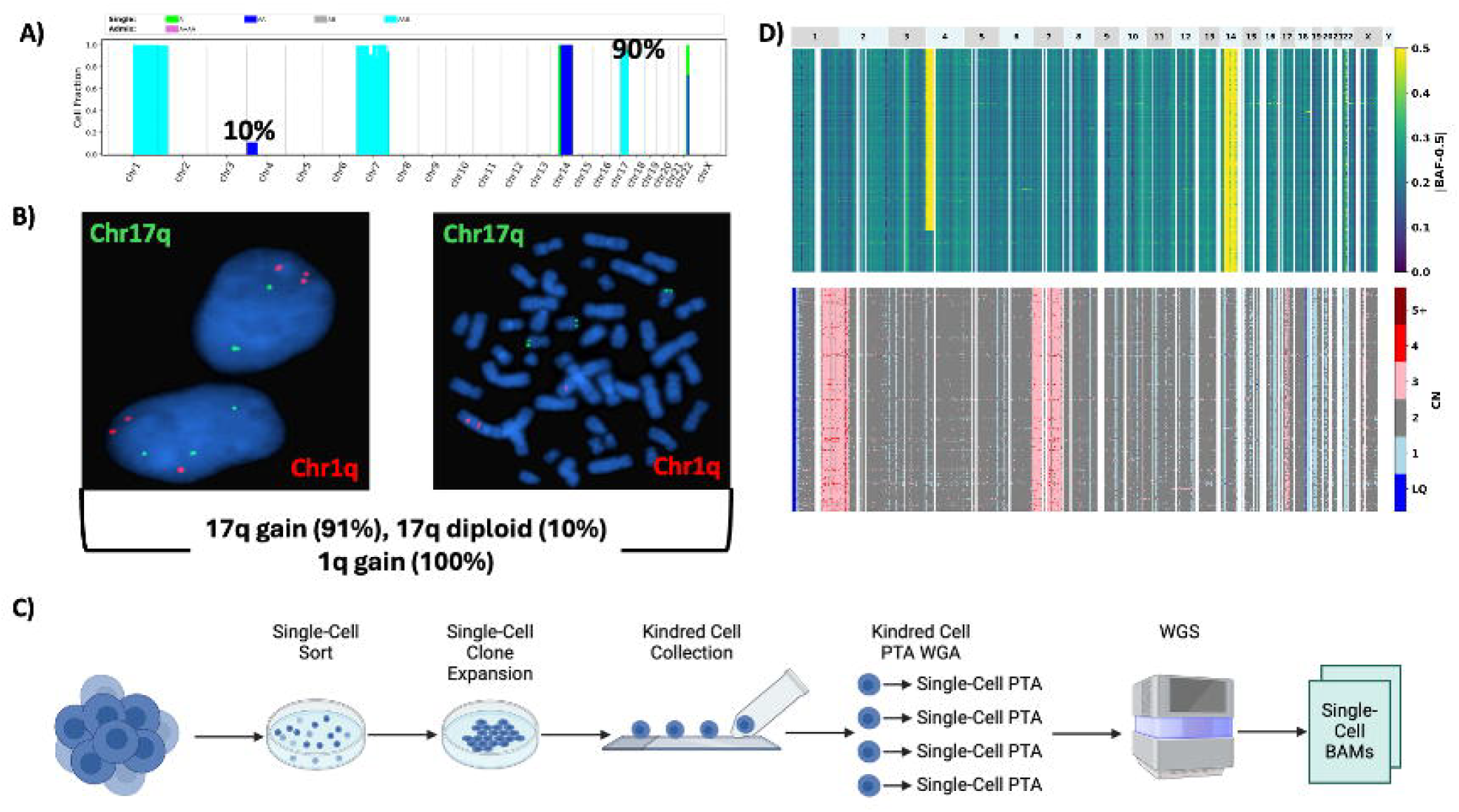

BE2C was derived from BE2, which was established from the bone marrow of a child with malignant neuroblastoma. S2K revealed different ploidy in these lines: hypodiploid for the parental line BE2 (**Fig. 5A**) and an admixture of hypodiploid and hypotetraploid in BE2C (**Fig. 5B**). These were validated by cytogenetic profiling of 100 metaphases in each line (**Fig. 5C, D**). In BE2, S2K showed a discrepancy between inferred copy number and allelic imbalance (**Fig. 5A**). While estimated copy number indicates a clonal 1-copy loss of chr18 (validated by FISH, **Fig. 5E**), the allelic imbalance value of 0.19 was far below the expected 0.5 from such loss (**Supplementary Fig. S6A**). Therefore, we hypothesized that the apparent discrepancy was caused by mirrored CNVs, i.e. loss of the different parental copy of chr18 in different subclones. To test this hypothesis, we performed droplet-based scRNA-seq (10X Genomics, **Fig. 5F**) to reduce the potential selection bias in plate-based single-cell profiling. Indeed, 1-copy loss of chr18 was present in nearly all 6,593 cells that passed QC checks with a major and minor subclone resulting from loss of different chr18 haplotypes detected at paternal haplotype frequency (pHF) of 0.81 and 0.16, respectively (**Fig. 5G left, Supplementary Fig. S6B**). The minor haplotype loss is significantly enriched in cluster 7 amongst the 9 clusters identified in scRNA-seq data (**Fig. 5G right**), which is retained in BE2C (with complete LOH, **Supplementary Fig. S6C**). These data support convergent evolution of chr18 loss in BE2 leading to two lineages retaining different haplotypes, and the one with the minor haplotype seeded BE2C which subsequently underwent WGD (**Fig. 5H**).

**Figure 5.**
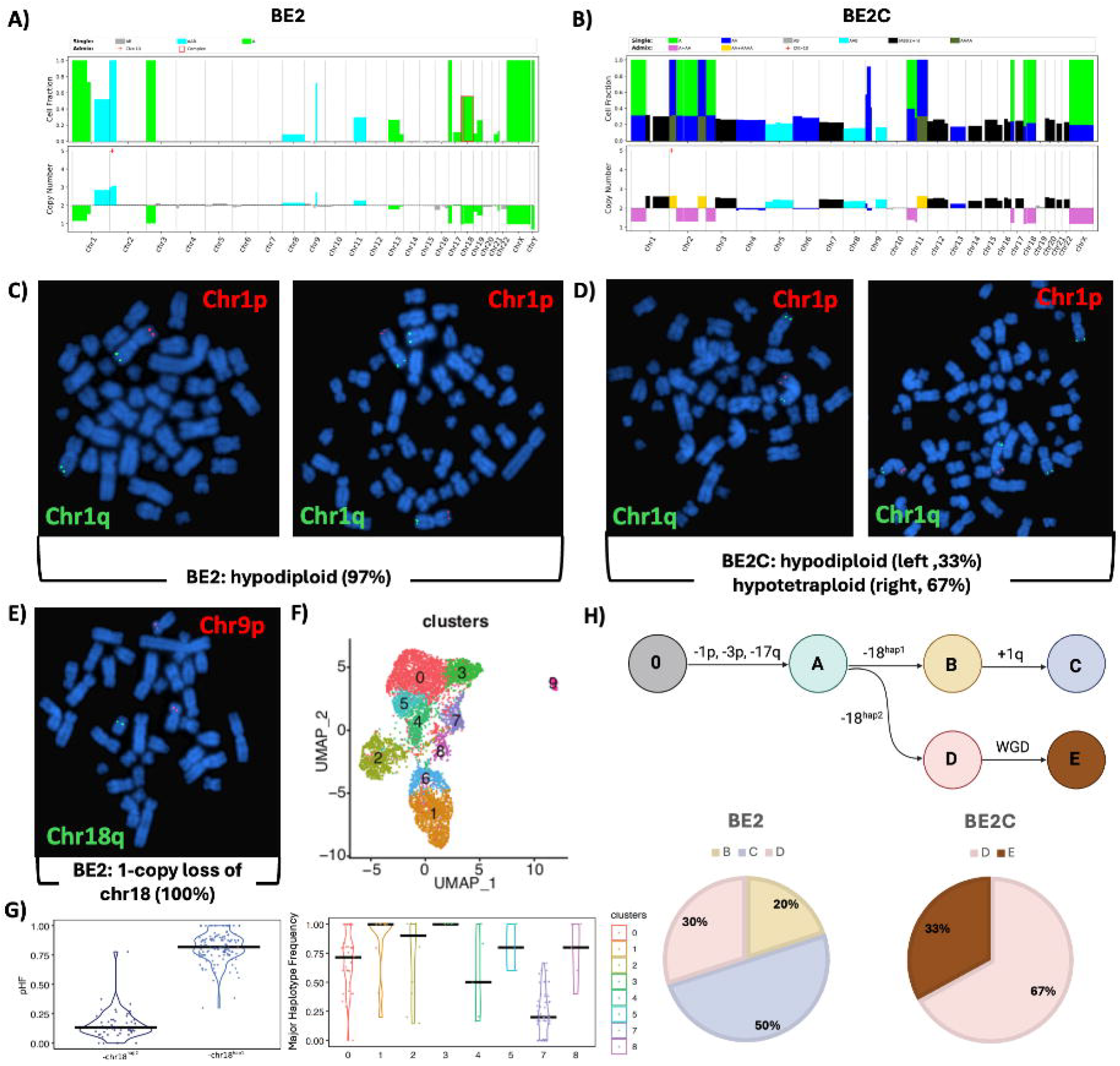

### 5. Patient sample analysis

To explore its potential use in clinical settings, we applied S2K to 23 AML and their matched germline samples, each with clinical annotations including cytogenetic profiling of 20 metaphases. The results, summarized in **Supplementary Table S4**, showed that S2K identified 47 out of the 51 abnormal cytogenetic events documented in the clinical reports with an additional 21 abnormalities detected exclusively by S2K. Interestingly, CNV analysis by DRAGEN matched S2K’s results on all discrepancies, suggesting that the differences may reflect ITH. The analysis on the 23 germline samples showed 1-copy gain on chromosome 21 for two cases with Down Syndrome.

The second use case involved the analysis of metastatic neuroblastomas with more complex CNVs than AML. The high concordance between published tumor purity and that estimated from clonal CNVs by S2K of this cohort (Pearson’s correlation coefficient = 0.95, *p* < 1E-22, **Supplementary Fig. S3C**) demonstrated an overall high accuracy of S2K. Similar to neuroblastoma cell lines (**Fig. 5B**), admixture of tetraploid and diploid cells was also found in several patient samples (**Supplementary Fig. S7**). To illustrate how subclonal CNVs can inform our understanding of tumor evolution, we reconstructed the CNV-based phylogenies for H134722, a case with multi-region sequencing data noted to have an inconsistent evolution trajectory projected by CNV versus point mutations in the original publication [19]. Using five WGS samples obtained from spatially distinct regions, S2K revealed both clonal and subclonal CNVs with varying CCF across the samples (labeled R3-R7 in the original study, **Fig. 6A**). A CNV-based evolution tree constructed with the parsimony principle [20] projected seven events (including MYCN amplification) detected in all samples as ancestral (**Fig. 6B**), including two events, gain of chr2 and chr7, that were subclonal in a subset of the samples. Under this scenario, the subclonal chr7 gain in R6, which had a CCF of 0.2 and shared the same amplified haplotype as the other samples (**Supplementary Fig. S8)**, would have been considered the initiation event with maximum parsimony. However, R6 was unlikely to seed the metastatic spread given its distance (kidney) from the sites where a prior biopsy and other metastatic samples were collected [19]. Therefore, the chr7 event in R6 was modeled as a subsequent subclonal loss to the initial copy gain (reversion) instead. Similarly, reversion of chr7 gain may have also occurred in R7 which was projected to have an admixture of cn-LOH and 1-copy gain of chr7. The resulting CNV-based phylogenetic tree is comparable to that constructed from point mutations, and the inferred clonal composition of each tumor region is consistent with the published model of metastatic spread after therapy.

**Figure 6.**
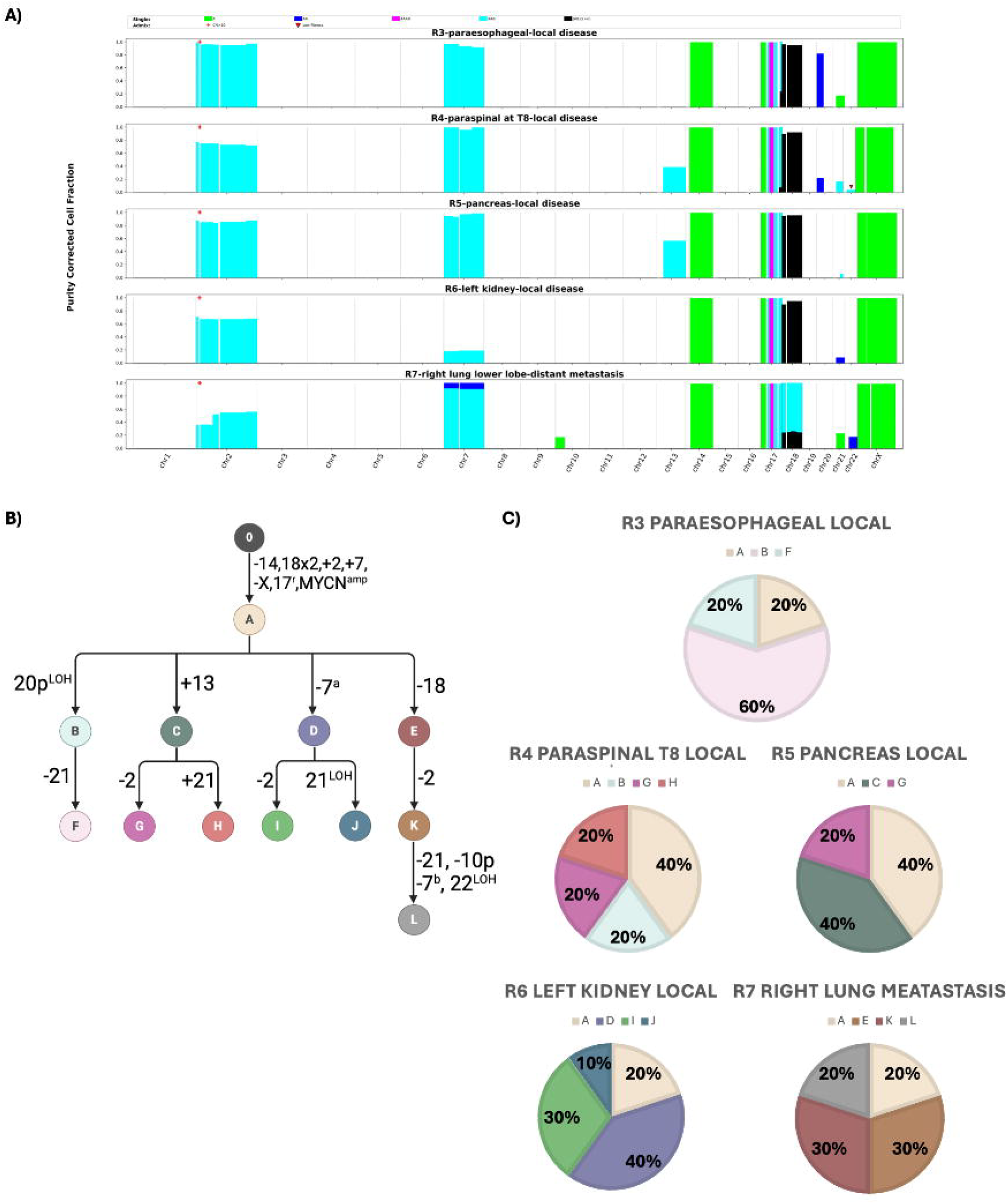

## Discussion

The Seq2Karyotype (S2K) algorithm, designed to perform digital karyotyping for single-sample WGS data, demonstrates a significant advancement in the analysis of CNVs and clonal heterogeneity. Although detection of subclonal CNV in a bulk sample has been implemented in existing tools [6-9], S2K is unique in modeling admixed CNV states, thereby enhancing our understanding of ITH and evolution as demonstrated in the diverse cohorts analyzed here. Furthermore, support for single-sample analysis is particularly valuable for analyzing tumor samples that lack matched normal samples, such as cancer cell lines or tumor sequencing for clinical oncology testing. When the results are integrated with SV breakpoint analysis, derivative chromosomes can be constructed (**Figs. 2 and 3**) which can further enhance the resolution of karyotype predictions. Single-sample analysis also makes S2K a useful method for analyzing disease-relevant germline CNV events—we performed an exploratory S2K analysis which detected disease-relevant mosaic uniparental disomy of 6p and 7p in patients with myeloid neoplasms (**Supplementary Fig. S9**), the later was also validated by scWGS (data not shown).

While CNV-based ITH has recently come to light for the two well-studied adult cancer benchmark cell lines, COLO829 and HCC1395, based on scWGS and cytogenetic profiling, detection of newly acquired subclonal CNV events in HCC1395BL, the matching germline for HCC1395, was unexpected. Furthermore, presence of the extensive ITH, including subclonal tetrapolidy, in 16/17 neuroblastoma cell lines profiled (**Supplementary Table S2**), indicates that CNV ITH is likely a common phenomenon in cancer cell line models. Extensive experimental validation, including SKY mapping, FISH and scWGS or scRNA-seq, corroborated S2K’s inference of CNV ITH in these cell line models. The evolving CNV profiles of these cancer models may pose challenges in using these models for benchmark analyses, and warrants future research to examine their effect on conferring selective therapeutic resistance, which was reported previously in tetraploid cells against DNA-damaging agents [21].

The evolving nature of CNV-based ITH in both cancer cell lines and patient samples emphasizes the importance of integrating CNV data with point mutations in evaluating the trajectory of tumor clonal evolution. This may involve consideration of convergent evolution such as the mirrored chr18 deletion in the BE2 cell line (**Fig. 5**) or reversion evolution such as the loss of der18 in COLO829 (**Fig. 2**). Reversion evolution should also be examined for multi-region tumor evolution analysis, as we showed that this could result in a phylogenetic tree more consistent with point mutation-based models and the known metastatic timeline than a tree constructed by the parsimony principle (**Fig. 6**).

S2K has several limitations. While the use of SNP-based data simplifies the preparation of input data, it may limit S2K’s performance in regions with low SNP density or high sequencing noise. Additionally, while S2K is optimized for large-scale CNVs, balanced translocation, reportable by cytogenetic profiling, requires analysis of structural variations using other available methods [22-24]. Also, to avoid overfitting, the current implementation only considers a maximum of two CNV states with the same direction (i.e. both events are copy gain or copy loss which may include LOH events), which may lead to loss of complexity in some regions. For example, our model projected that 50% of the NB5 cells contain 1-copy loss of chr3 except for a 21Mb segment at 3qter which had 1-copy gain in 50% cells. The SKY mapping indicates that the elevated copy-number at this segment was caused by a clonal copy number gain involving interchromosomal translocation which was followed by a subsequent 1-copy loss of the entire chromosome 3 in a subset of cells (**Supplementary Fig. S10)**. Such a region with copy gain and loss occurring simultaneously in the same cell is not captured by our model.

In conclusion, S2K is a powerful tool for analyzing genomic heterogeneity and clonal architecture by deconvoluting CNV events in bulk tumor samples. By enabling in-silico karyotyping at high accuracy for individual tumor samples, we demonstrated S2K’s utility for both research and clinical applications. The unexpected intratumoral heterogeneity of CNV events detected in cancer cell line models by S2K analysis highlights a need to understand tumor evolution and therapeutic resistance using these models for future research.

## Materials and Methods

### 1. Data sets and validation

| Dataset | Description | Number of Samples | Purpose |
| --- | --- | --- | --- |
| **Adult Cell Lines** | COLO829 and HCC1395 with matching germline samples (COLO829BL, HCC1395BL) | **4** | Benchmarks for validating the S2K pipeline using well-characterized genomic profiles. |
| **Neuroblastoma Cell Lines** | NBL cell lines with high genomic instability, complex karyotypes, and extensive clonal heterogeneity. | **17** | Evaluates S2K's ability to detect major and minor subpopulations and karyotypes in genomically unstable cell lines. |
| **AML Patient Samples** | AML patient samples with matching germline samples. | **46** | Tests S2K's utility in analyzing hematologic malignancies, assessing its accuracy in identifying CNVs and clonal heterogeneity against clinical standards. |
| **CCDI Dataset** | Neuroblastoma diagnostic samples from 142 subjects [19]. | **184** | Challenges S2K's capability to resolve tumor subpopulations in complex NBL genomes, considering variability in clonal architecture and tumor purity. |

### 2. S2K Algorithm

#### 1. Coverage and B-allele fraction calculation using SNP allele counts

The input for S2K is a user-provided file containing essential information about SNPs identified from WGS data. This file includes the chromosome and position of each SNP, along with the read counts for both the reference allele (A allele) and the variant allele (B allele), labeled as *n*_*A*_ and *n*_*B*_, respectively. Only SNPs that match a high-quality (HQ) SNP list are used for calculating coverage *m*(*m*= *n*_*A*_ + *n*_*B*_) and B-allele fraction 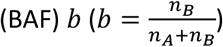. While S2K provides a default HQ SNP list (**Supplementary Information**), users may substitute their own SNP list—for example, one generated from a matched normal sample. This ensures the robustness of the mand b values. SNPs projected to have heterozygous genotypes are further analyzed to identify patterns and clonalities of copy number variations (CNVs) and copy-neutral loss of heterozygosity (cn-LOH). These changes are expected to cause deviations from the expected BAF of 0.5 in a diploid region, which can also be accompanied by changes in coverage in the case of CNVs.

#### 2. Diploid chromosome detection

Diploid chromosomes, which serve as the normal reference, are identified through a two-step analysis. In the first step, S2K examines the joint distribution of BAF (b) and coverage (*m*) of all SNPs within a chromosome. X and Y chromosomes are excluded from this analysis. The joint distribution is described as follows:

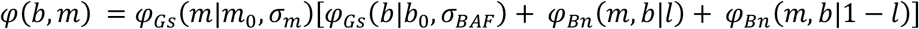

where *φ* denotes the joint probability density function of BAF and coverage while *Bn* and *Gs* denotes Binomial and Gaussian distribution, respectively. *l* represents the noise (i.e., sequencing error) in measuring homozygous A and B genotypes as shown in the second and third terms, respectively. Binomial and Gaussian are used to ensure the robustness of modeling.

The width of BAF distribution, denoted *σ*_*BAF*,_ is estimated by a binomial approximation as follows:

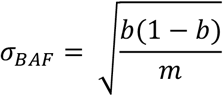

The width of the coverage distribution is estimated by Poisson approximation. To account for deviation from the model in the observed data, the standard deviation of coverage is adjusted by a factor *w*, such that *w* > 1 *and w* ≈ 1:

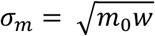

The parameters of the *φ* distribution, coverage, BAF, and factor w, are determined by minimizing the following χ^2^:

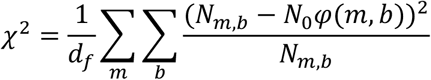

Where *d*_*f*_ is the number of degrees of freedom. This double sum enumerates all possible pairs of coverage and BAF within the range of sample coverage (1^st^ to 99^th^ percentile by default). It provides a normalized difference between the observed and modeled 2-D histogram of (b, *m*).

In the second step, chromosomes with prominent CNVs (e.g., clonality ≥30%) are expected to have outlier parameters. This is analyzed using a generalized Extreme Studentized Deviate (ESD) many-outlier procedure [25], with a slight modification: the outlier threshold is estimated based on a normal distribution rather than by selecting a subset of data points to avoid boundary problems. If the ESD procedure fails, percentiles of the parameters are used to evaluate outlier status. Chromosomes identified as outliers are considered non-diploid by this step, while those deemed diploid will serve as the initial normal diploid reference for subsequent analysis.

#### 3. Initial segmentation and labeling

All chromosomes are initially segmented into one million base pair windows. Each segment is evaluated by S2K to determine the ploidy (diploid, non-diploid, unknown) by examining the SNPs within corresponding segments. First, segments with CNVs at high clonality (≥30%) are analyzed using the parameters derived from the previous step. In each segment, SNPs are labeled as nondiploid (E) or diploid (D) based on the assessment of a random variable *z*, which is expected to obey *χ*^2^ distribution with two degrees of freedom:

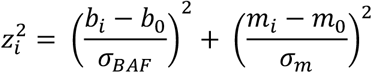

Note that for diploid and non-diploid chromosomes derived from the previous step, the parameters for reference (i.e., *m*_0_ and b_0_) are based on the native chromosome and genome-wide average, respectively. Next, S2K performs evaluation of SNPs in segments with low clonality, which are deemed as putative diploid regions. For each segment, the BAF distribution of all D-label SNPs is modeled by the binomial distribution shown below and is required to be convoluted with distribution of coverage:

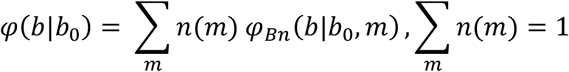

Parameters are estimated by minimizing χ^2^ value which is expected to be 1 for diploid segments; a higher χ^2^ value reflects the degree of non-diploid state. Furthermore, a non-diploid segment can result in outlier values of at least one estimated parameter, which is identified using the same outlier detection approach described in the preceding section. To define such regions, a sliding window encompassing 1,000 SNPs (by default) is analyzed by assessing whether allelic imbalance (*ai*) deviates from the diploid. A segment with significant deviation is subjected to a localized re-analysis of BAF and coverage to verify its outlier status and once confirmed, SNPs within such segments are relabeled as unknown (U). After all analysis is completed, the ploidy (i.e., label) of the segment is assigned based on the majority label of the SNPs within the segment.

#### 4. Segment refinement

After the initial segmentation and parameter estimation, the segmentation refinement process is carried out using a detailed two-step procedure.

In the first step, S2K employs a change-point-detection (CPD) algorithm [26] to refine the segments. This algorithm identifies change points for each chromosome based on the segments’ allelic imbalance (*ai*) and coverage. Since the number of change points is not known beforehand, the penalized optimization balances the fit of the model with the complexity introduced by additional change points. This approach ensures that the segmentation is neither over-segmented nor under-segmented, achieving an optimal balance. Once the change points are identified, the first round of segment refinement is executed by merging segments between these change points. The goal here is to provide a broad refinement that captures the major structural changes in the genomic data.

After the initial refinement using the CPD algorithm, S2K proceeds with a detailed recursive merging process. In this step, S2K re-estimates the allelic imbalance (*ai*) and coverage parameters for the newly refined segments. It then evaluates whether neighboring segments should be merged based on these parameters. Specifically, if the *ai* or coverage of two neighboring segments differ by more than a user-defined threshold, the segments remain separate. Conversely, if the parameters are within the threshold, the segments are merged to form a more homogeneous region. This merging process is carried out recursively, with S2K continuously re-evaluating and merging segments until no further segments meet the criteria for merging. This iterative approach ensures that the final segmentation is finely tuned, accurately reflecting the underlying genomic structure and enhancing the precision of subsequent analysis.

#### 5. Karyotype modeling

With the refined segments in place, the next step involves karyotype modeling to further analyze the genomic structure and variations. The distribution of coverage (*m*) and allelic imbalance (*ai*) in non-diploid segments is modeled to identify the most fitting CNV pattern (i.e., karyotype) and clonality (i.e., cellular fraction). Adopting the convention of using A and B to denote the two haplotypes in a diploid region, the evaluated models include: AAB for one-copy gain, A for one-copy loss, AA for cn-LOH, (AB)(2+n) for bi-allelic duplication, and (AB)(2-n) for bi-allelic loss, among others. Additionally, many complex models involving two karyotypes, such as A+AA, AA+AAB, etc., are also evaluated. Below are example model functions for coverage (*m*) and allelic imbalance (*ai*), with clonality (denoted as *k*) as a variable. The detailed list of functions are presented in **Supplementary Information**.

For one-copy loss (A): coverage *m*is calculated by averaging number of alleles, as follows:

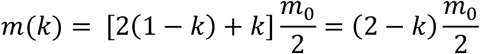

where *m*_0_ denotes the coverage of the diploid regions

BAF and allelic imbalance (*ai*) can be calculated below:

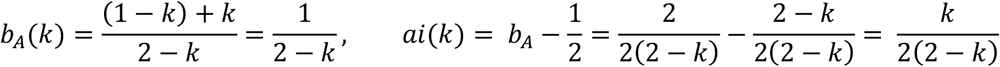

For one-copy gain (AAB):

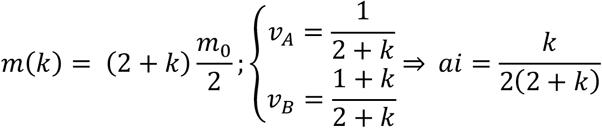

For cn-LOH (AA):

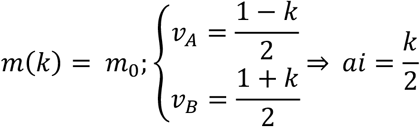

For bi-allelic gain (AB)(2+n) or loss (AB)(2-n), only coverage change is considered due to lack of allelic imbalance:

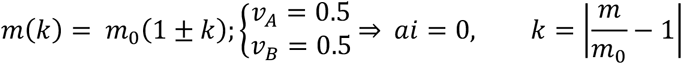

Other models can be derived following the same logic. Note for complex models, the clonality *k* denotes the clonality of the latter component.

With the modeling for each karyotype complete, we proceed to select the appropriate model for each segment. This is achieved by assessing the alignment between theoretical values and observed data. This alignment, or fitness, of each model is quantified by calculating a customized Euclidean distance between the theoretical values and the observed measurements. Given the estimated clonality (*k*) for a given model (*M*) and segment (*S*), the distance can be calculated as follows:

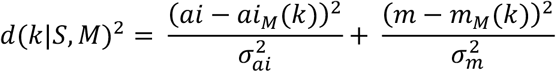

The model (*M*) with lowest distance will be assigned to segment (*S*). However, this can lead to a problem of severe overfitting, where S2K may select overly complex models. In order to combat this issue, we introduced a penalty term (*τ*) for each model based on the model evolution. For example, the penalty for one-copy loss (A) is 2, given that two steps are required to achieve such a state, namely lose one copy then estimate the clonality. The penalty term is calculated for all models available. For a detailed penalty list, please refer to supplementary information. Thus, the final model selection can be defined as:

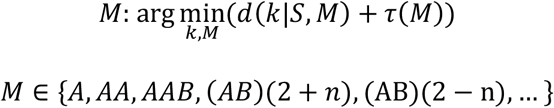

Since the equations are non-linear, the distance is determined numerically.

#### 6. Karyotype refinement

To enhance the accuracy of karyotype modeling, S2K uses a Gaussian mixture clustering approach [27] to refine karyotype assignments. This probabilistic method groups genomic segments based on similarities in copy number and allelic imbalance, enabling more precise characterization of underlying CNVs. By clustering segments with comparable profiles, the method ensures consistent treatment of similar patterns, reducing noise and improving karyotyping reliability.

For each cluster, the most likely karyotype model is selected by calculating the distances between the observed data and all potential models. The model with the smallest distance is chosen, ensuring consistent clonality across clustered segments and providing an accurate depiction of the genomic landscape.

#### 7. Visualization

Visualization plays a crucial role in S2K, providing insights into CNVs and facilitating the tuning of reference diploid and merging parameters. Segments assigned to karyotype models are displayed through various views, including a genome-wide overview (**Fig. 2A**) and chromosomal views showing BAF, coverage, segment boundaries, and projected clonality. By default, the genome-wide overview displays segments larger than 5 Mb that overlap less than 50% of the centromere, while the chromosomal view includes segments over 1 Mb with the same centromere overlap criteria. These thresholds are user-adjustable. Users can evaluate the fit of allelic imbalance and coverage to assigned models, allowing reassessment of reference diploid selection, as illustrated in **Supplementary Fig. S1D**.

#### 8. Tumor purity estimation

The cancer cell fraction (CCF), or tumor purity, can be estimated from S2K results. To calculate CCF, kernel density estimation (KDE) is applied to the clonality distribution of non-diploid segments, allowing systematic identification of clones based on peak prominence in the KDE plot. Local maxima in the plot represent distinct clone groups. The major clone is determined by the most prominent peak—defined as at least 40% higher than the others—or, if peak prominence is similar, the peak with the highest clonality. This quantitative method ensures a reproducible approach for defining the major clone and calculating S2K-based CCF.

### 3. Data availability

**COLO829**: The matched tumor/normal whole-genome sequencing (WGS) dataset for the COLO829 melanoma cell line is available under dbGaP accession **phs000932.v1.p1**.

**HCC1395**: WGS data for HCC1395BL cell line is available in the NCBI SRA under accession **PRJNA1279751. Neuroblastoma Cell Lines**: WGS data for all 17 neuroblastoma cell lines are available in the NCBI SRA under accession **PRJNA1279751**.

**AML Patient Samples**: Sequencing data for all AML patient samples are available through **St. Jude Cloud** (https://platform.stjude.cloud/data/cohorts/pediatric-cancer), with corresponding IDs listed **in Supplementary Table S4**.

**CCDI Neuroblastoma Study**: Whole-genome and transcriptome sequencing data from the CCDI neuroblastoma study are available under dbGaP accession **phs003111.v1.p1**.

### 4. Validation experiments

#### Cytogenetic experiment

Cytogenetic analyses such as fluorescence in situ hybridization (FISH) and spectral karyotyping (SKY) are integral components of genomic characterization in both clinical and research settings. These assays are conducted by the Cytogenetic Shared Resource Laboratory at St. Jude Children’s Research Hospital using standardized protocols. FISH involves labeling purified bacterial artificial chromosome (BAC) DNA probes (Supplementary Table S3) with fluorescent dyes via nick translation, followed by hybridization to metaphase chromosomes or interphase nuclei. After overnight incubation and post-hybridization washes, nuclei are counterstained with DAPI and scored under fluorescence microscopy to determine copy number or detect rearrangements at targeted loci. SKY analysis begins with metaphase chromosome preparation and trypsin-Giemsa banding, followed by hybridization with a combinatorially labeled chromosome paint probe mix. Spectral imaging and computer-assisted classification assign pseudocolors to each chromosome, enabling the resolution of complex structural abnormalities and derivative chromosomes. These assays are interpreted and reported by board-certified cytogenetic technologists and cytogeneticists, ensuring analytical rigor and quality control. Together, FISH and SKY offer complementary insights into chromosomal architecture and clonal diversity, enhancing the resolution and confidence of genomic findings. The details on the DNA probes, number of interphase and metaphase cells profiled for FISH and SKY mapping are described in **Supplementary Table S3**.

#### Bulk sample whole genome sequencing of cancer cell lines

Genomic DNA was quantified using the Quant-iT PicoGreen dsDNA assay (Thermo Fisher Scientific). DNA fragmentation was performed using a Covaris LE220 ultrasonicator to achieve the desired fragment size. Library preparation was conducted with the KAPA HyperPrep Library Preparation Kit (Roche, PN 07962363001), following the manufacturer’s protocol. Library quality and insert size distribution were assessed using the Agilent 2100 BioAnalyzer High Sensitivity DNA Kit, the 4200 TapeStation D1000 ScreenTape assay, or the 5300 Fragment Analyzer NGS Fragment Kit (Agilent Technologies). Final libraries were quantified using the Quant-iT PicoGreen dsDNA assay. Sequencing was performed on an Illumina NovaSeq 6000 platform with 2 × 150 bp paired-end reads, targeting an average genome coverage of 60× per sample.

#### Single-cell whole genome sequencing

Live single cells (Calcein Positive) from the SY5Y cell line were directly sorted into 96-well plates and subjected to Primary Template-directed Amplification (PTA) according to the manufacturer’s protocol (BioSkryb ResolveDNA Whole Genome Amplification Kit, cat# PN 100068). The PTA process uses a proprietary termination base that produces a short fragment DNA pool with amplicons ranging from 250 to 1500 base pairs (bp) without biased amplification. Following PTA, the amplified DNA was purified using magnetic beads to remove amplicons smaller than 200 bp and was subjected to the KAPA Hyper Prep Library Preparation kit (KK8504). The library preparation started 200ng of PTA amplified DNA followed by end-repair and A-tailing, followed by adapter ligation and library amplification following the manufacturer’s protocol. Subsequently, the amplified libraries were used to generate paired-end sequencing reads (2 × 150 bp) using Illumina NovaSeq 6000 for 300 cells, achieving approximately 10× coverage per sample. The sequences were aligned to the human reference genome (NCBI build 38) using BWA.

#### scRNA-seq

SK-N-BE(2) cells were passed through a 15 µm pluriStrainer (pluriSelect Life Science) to obtain a high-quality single-cell suspension. Single-cell RNA-seq libraries were generated using the Chromium Single Cell 3’ Library & Gel Bead Kit v3 (10x Genomics), following the manufacturer’s protocol. Libraries were sequenced on an Illumina NextSeq 2000 platform to ensure sufficient depth for transcriptome profiling. Raw sequencing data were processed using Cell Ranger v7.1.0 (10x Genomics), aligned to the GRCh38 human reference genome to generate gene-barcode matrices for downstream analysis.

#### Comparison with AML clinical report

Clinical karyotyping was performed for 20 AML cells per subject. To compare S2K results with AML clinical reports, tumor and matched germline WGS samples were analyzed separately, and the resulting data were analyzed to identify CNVs, LOH, and estimated CCF. The results from tumor WGS samples were converted to cytogenetic bands using the mapping of genomic locations for cytogenetic bands. Germline samples were evaluated for chromosomal level abnormality, such as trisomy 21 in Down syndrome.

### 5. Other analyses

Reference SNP datasets, scWGS analysis, scRNA-seq analysis, SV analysis are described in Supplementary Information.

## Software availability

Seq2Karyotype is publicly available on Github: https://github.com/chenlab-sj/Seq2Karyotype

## Acknowledgements

This study was supported in part by a Cancer Center Support Grant (P30 CA21765) to St. Jude Children’s Research Hospital, a Supplement of NCI Childhood Cancer Data Initiative (CA021765-41S3) to JZ, R01 CA216391 to JZ, R01CA262790 and R01CA266600 to XC and the American Lebanese Syrian Associated Charities. We would like to thank Michael Edmonson for his assistance with proofreading the manuscript.

---

**Figure S1.**
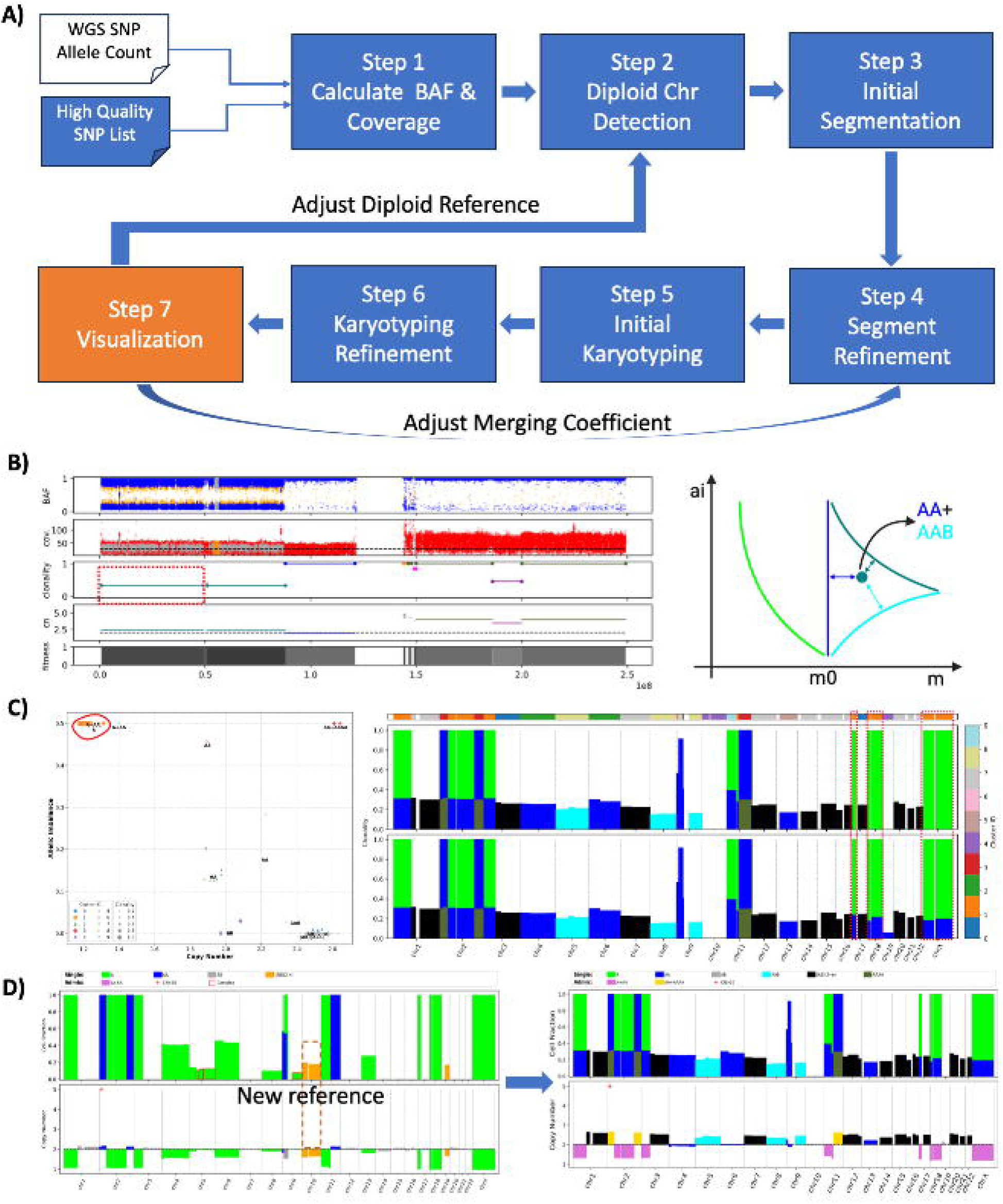

**Figure S2.**
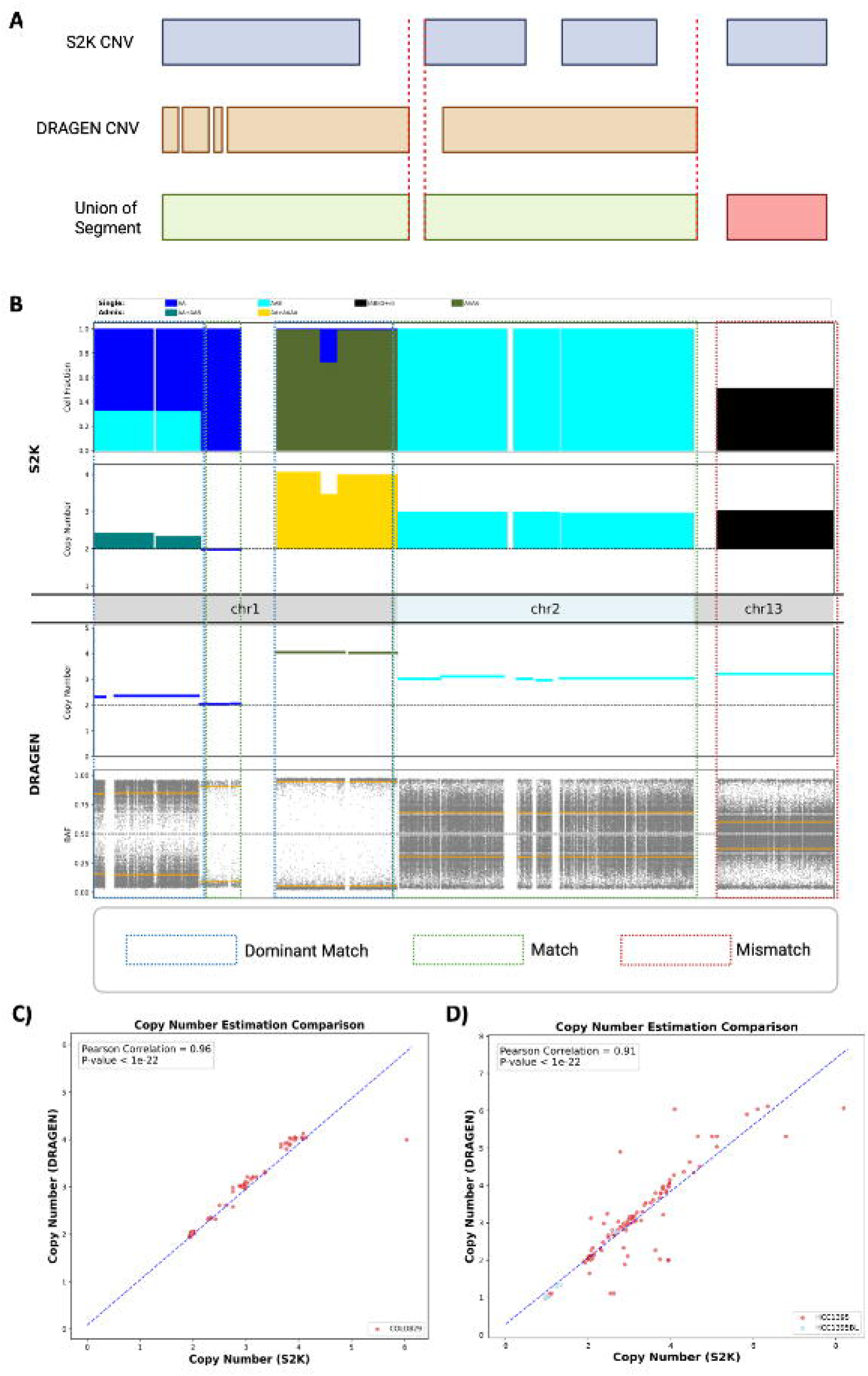

**Figure S3.**
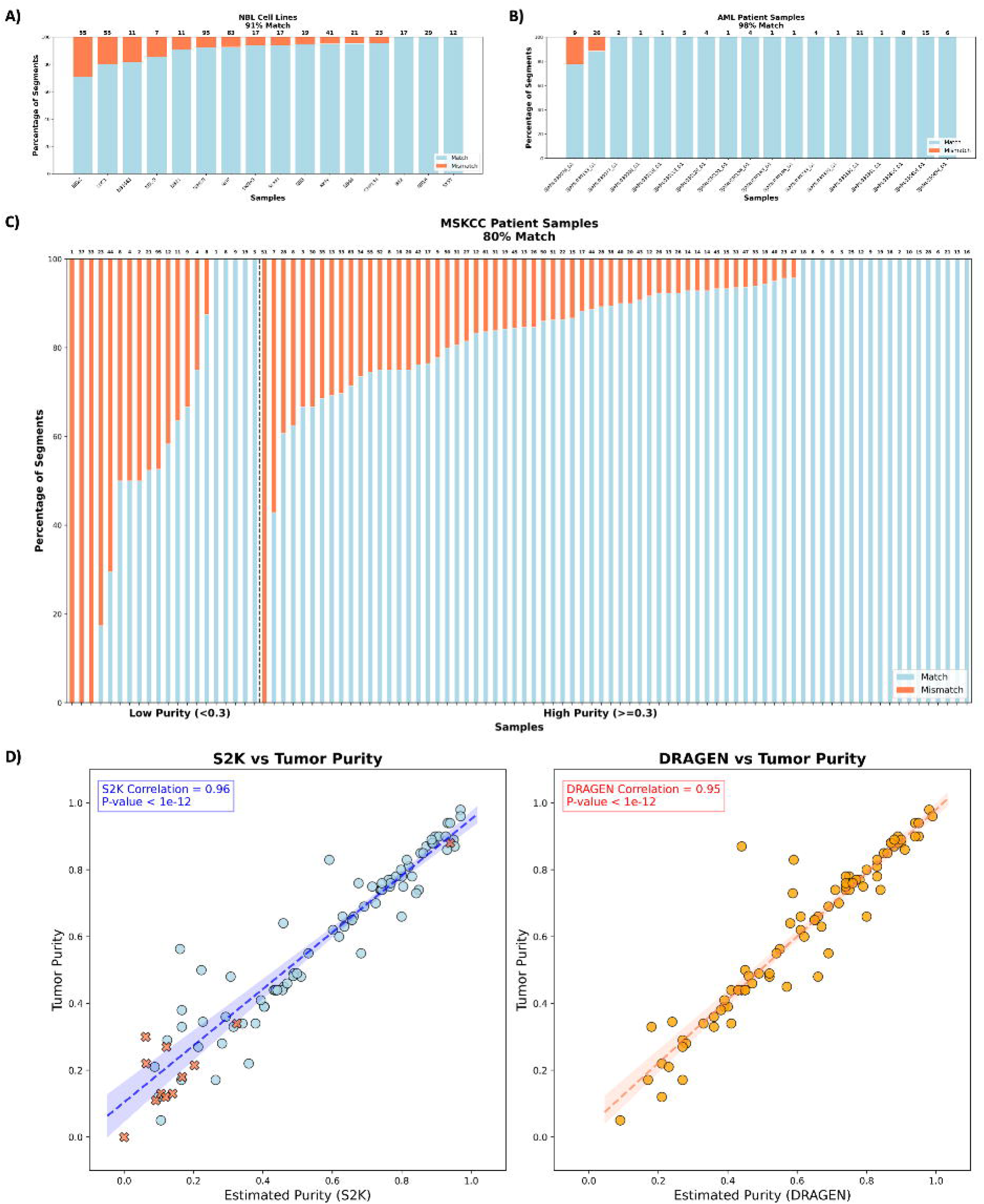

**Figure S4.**
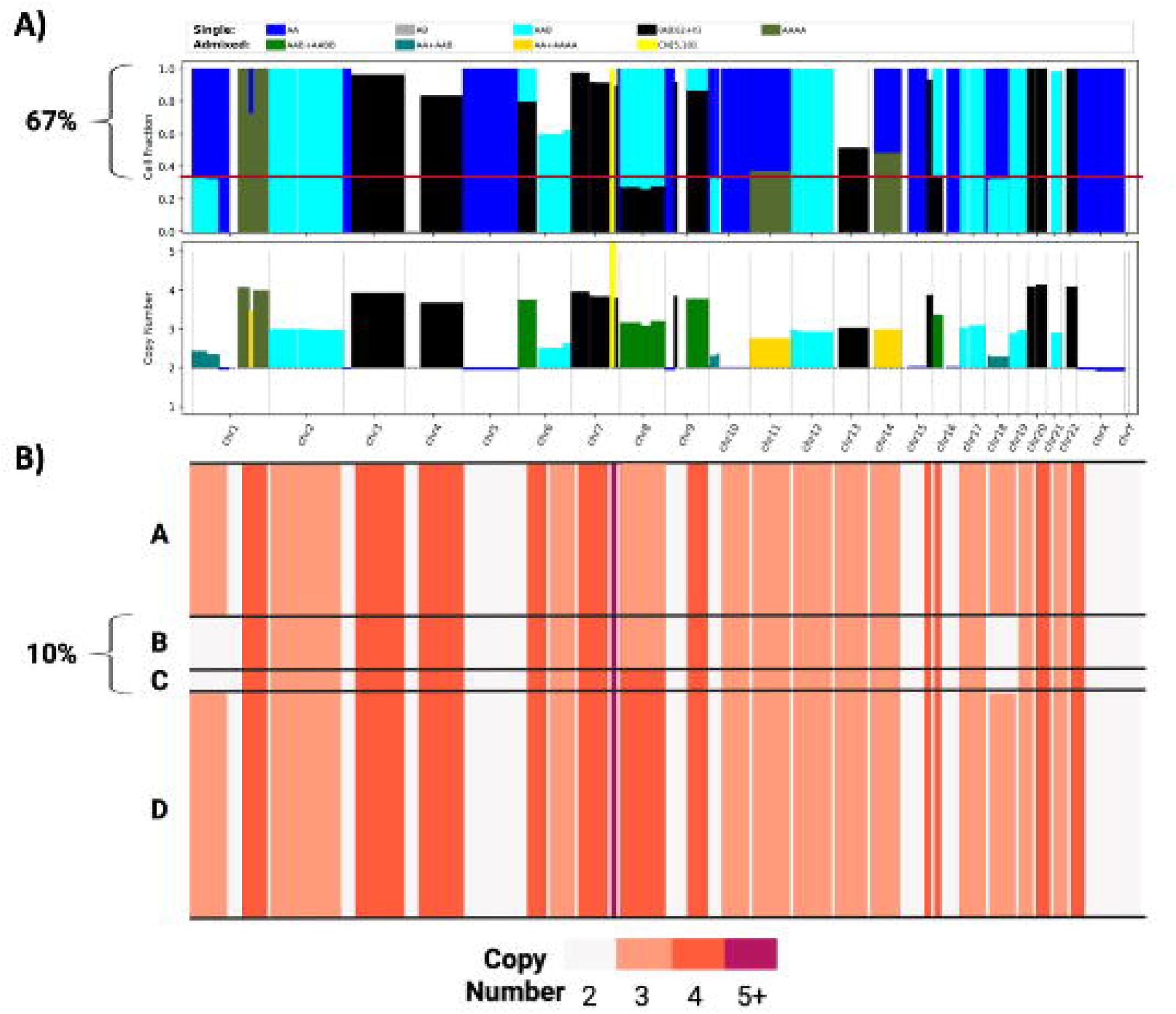

**Figure S5.**
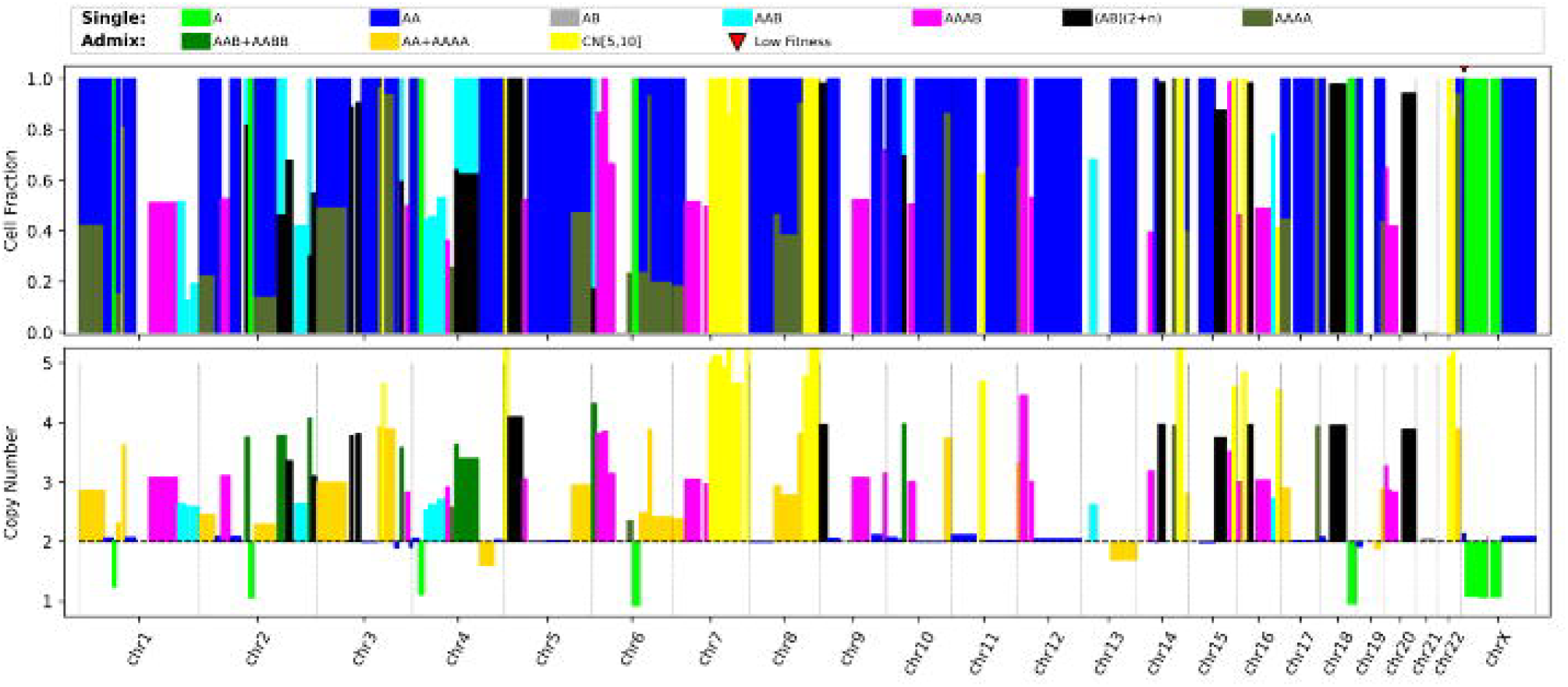

**Figure S6.**
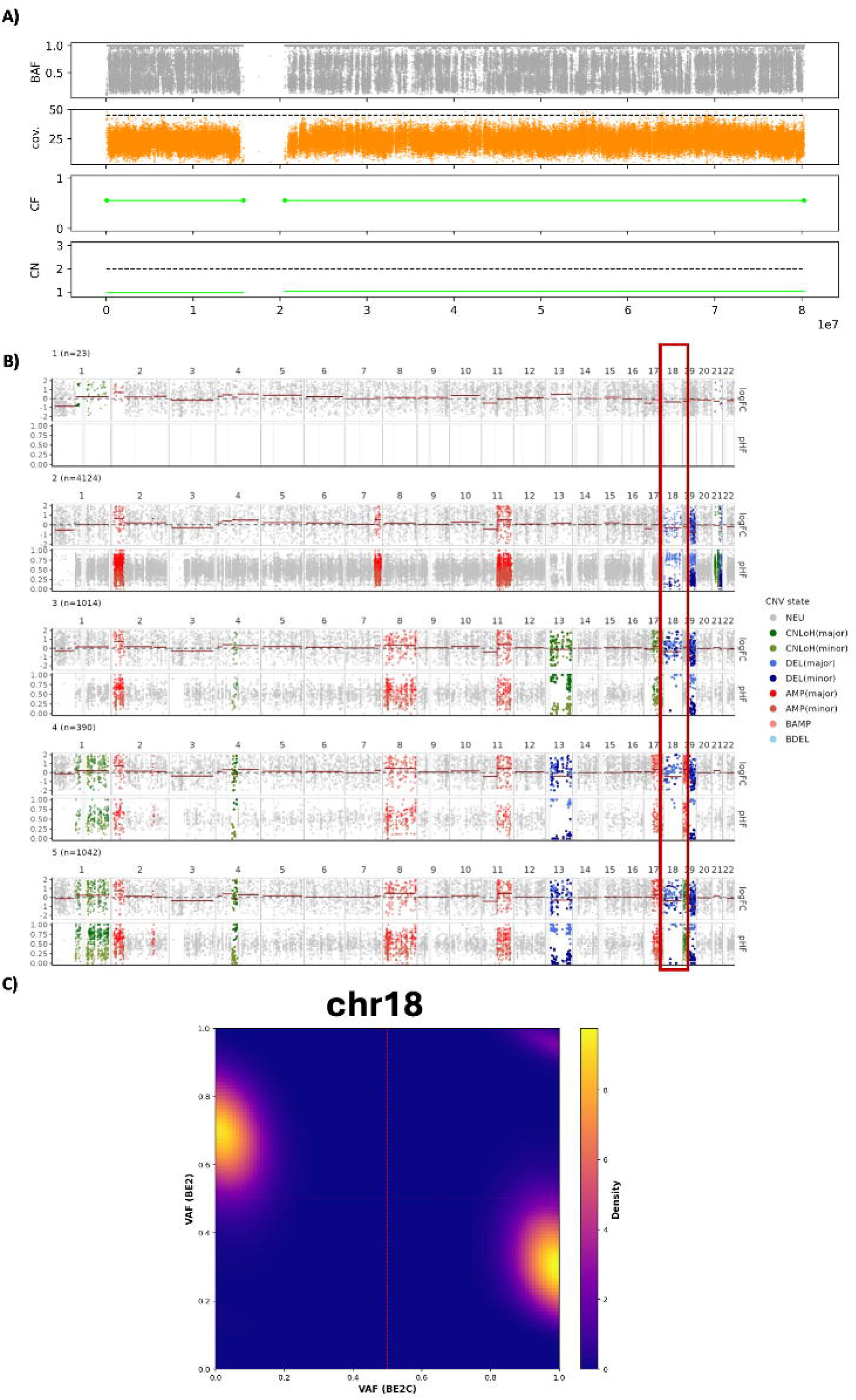

**Figure S7.**
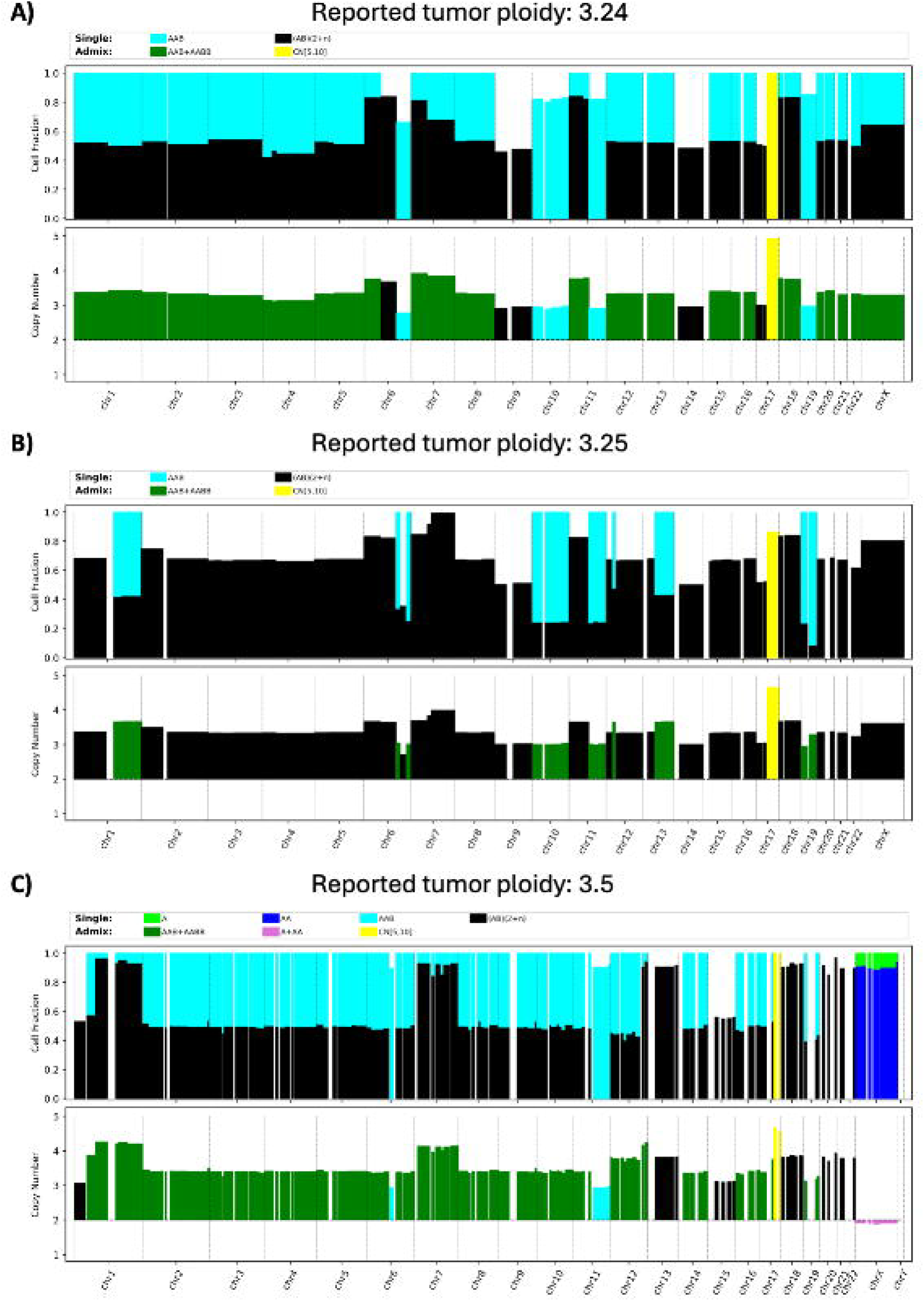

**Figure S8.**
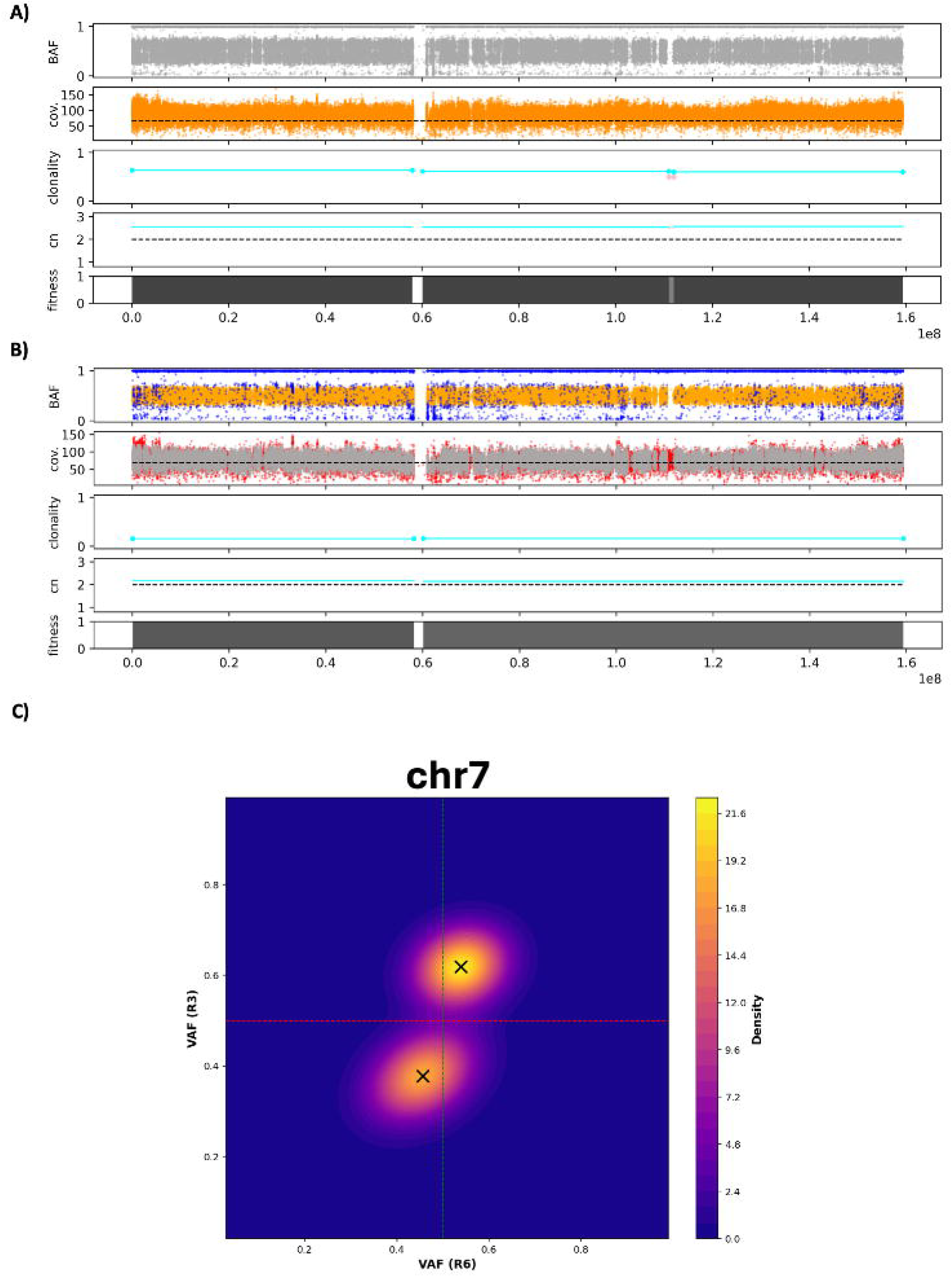

**Figure S9.**
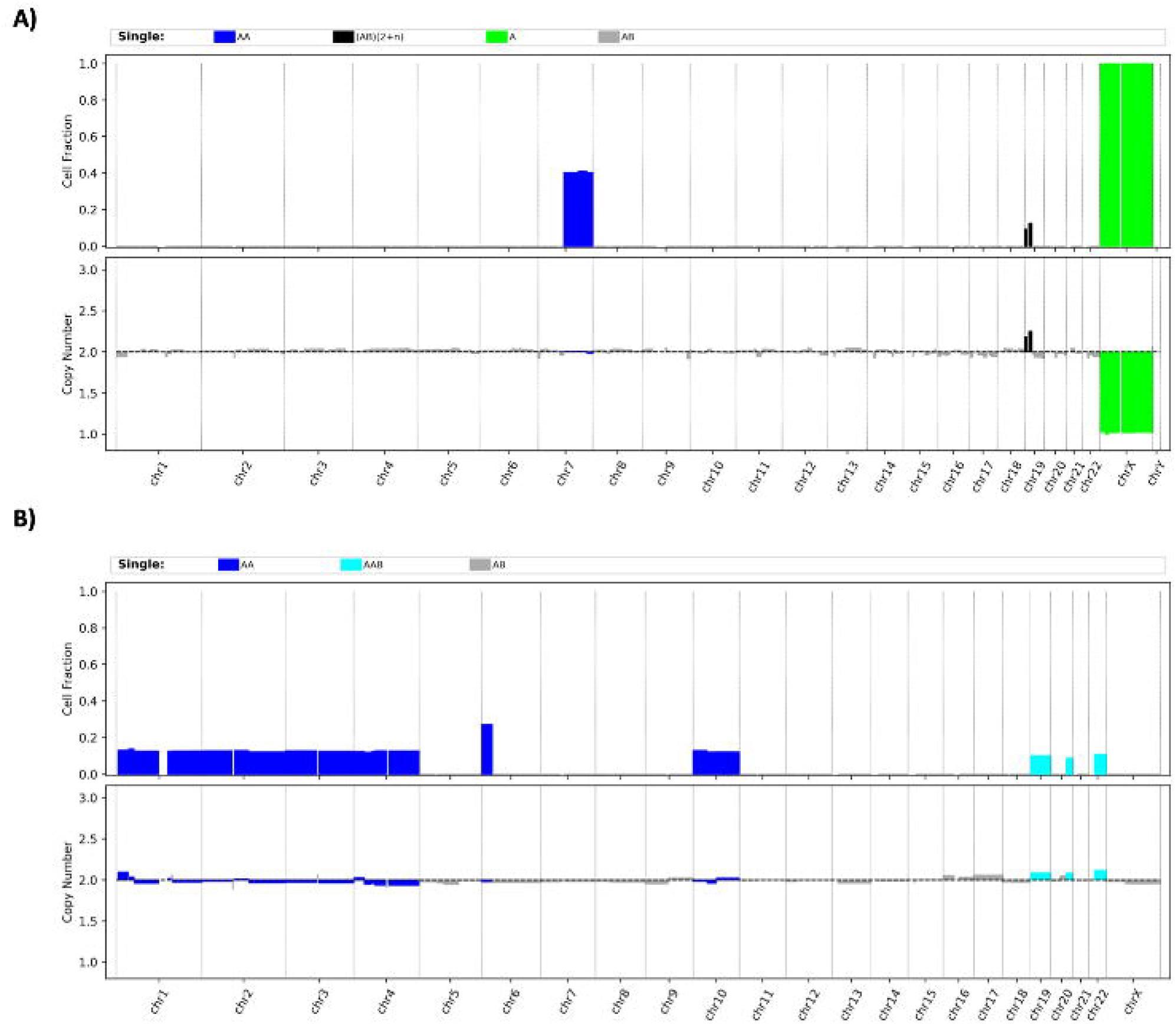

**Figure S10.**
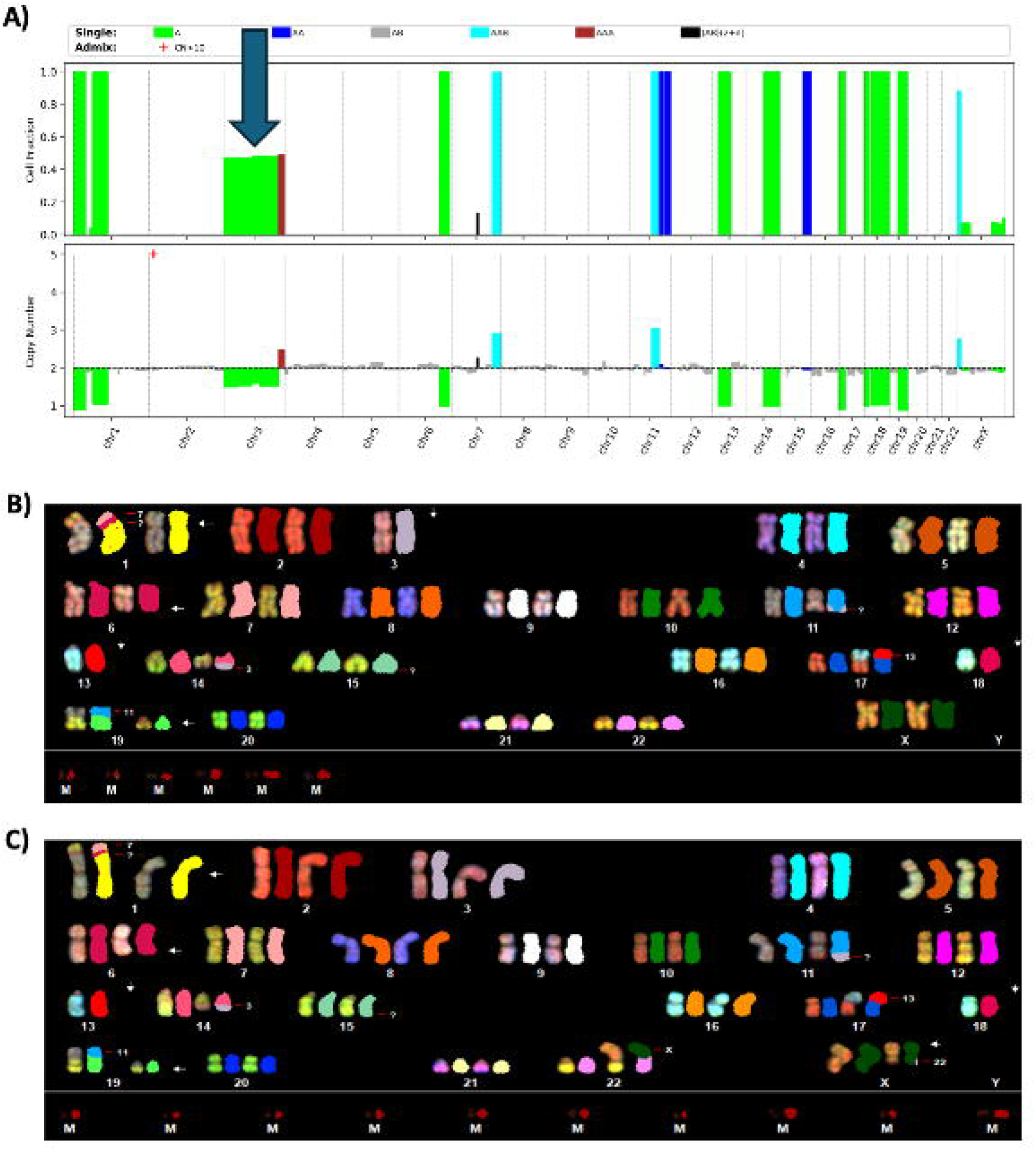

