## Supplementary information for "Seq2Karyotype (S2K): A Method for *in-silico* Karyotyping Using Single-Sample Whole-Genome Sequencing Data"

**Methods**

***S2K workflow***

S2K is a single-sample computational framework for inferring karyotype-level alterations from whole-genome sequencing (WGS) data. It requires as input a file listing the genomic coordinates and read counts of reference (A allele) and alternate (B allele) alleles for single nucleotide polymorphisms (SNPs). These SNPs are filtered against a curated panel of ~54 million high-confidence germline SNPs derived from >8,000 WGS samples to ensure reliability and facilitate normalization.

In **Step 1**, S2K computes two primary metrics—read coverage and B-allele fraction (BAF)—for each retained SNP. These distributions form the basis for identifying reference diploid chromosomes in **Step 2**, which serve as internal controls for subsequent modeling.

**Step 3** segments the genome into regions of homogeneous copy number and allelic imbalance based on the joint distribution of coverage and BAF. These initial breakpoints are refined in **Step 4** by merging adjacent segments with similar profiles and resolving borderline transitions.

In **Step 5**, each segment is evaluated against a compendium of candidate karyotype models, including single-copy gain or loss, bi-allelic gain, loss of heterozygosity (LOH), and complex CNV/LOH combinations. For each candidate model, S2K estimates the cellular fraction contributing to the signal, thereby capturing subclonal architecture within the tumor (Extended Data Fig. 1B).

To improve model consistency across the genome, **Step 6** clusters segments with similar copy number and allelic imbalance states and assigns a unified karyotype model to each cluster (Extended Data Fig. 1C). This step mitigates local noise and enhances interpretability in the presence of genomic complexity or subclonality.

Finally, **Step 7** incorporates an interactive visualization framework to support manual review and optional refinement of the segmentation and modeling results. This includes re-selection of reference diploid chromosomes and adjustment of segment boundaries, particularly in samples exhibiting complex aneuploidy or genome-wide duplication (Extended Data Fig. 1D).

Together, these steps enable robust and interpretable inference of large-scale somatic copy number alterations and their clonal structure from a single tumor genome, without requiring a matched normal control. The details of these steps are described in **Methods**.

***High-quality SNP list***

High-quality germline SNPs were identified from whole-genome sequencing data of 7,741 childhood cancer survivors (5,053 from the SJLIFE cohort and 2,688 from the CCSS cohort) hosted on Survivorship Portal (1) under accessions SJC-DS-1002 and SJC-DS-1005 on St Jude Cloud Genomics Platform (<https://platform.stjude.cloud/data/cohorts>). The variant data were processed using a four-step filtering pipeline based on genotype call rate, coverage distribution, variant allele frequency (VAF) distribution, and Hardy-Weinberg equilibrium (HWE). Variants were retained if they had a call rate ≥95%, non-outlier read coverage (Kolmogorov–Smirnov P > 1×10⁻²⁰), and VAF distributions consistent with expected germline genotypes (χ² < 22 for SNVs and < 67 for INDELs; composite P > 0.15 for SNVs and > 0.1 for INDELs). Variants also had to conform to HWE (P > 1×10⁻⁶). Filtering thresholds were optimized using ROC analysis against validated SNPs from GIAB, HapMap, and the 1000 Genomes Project.

***Karyotype function***

The table below summarizes the detailed karyotype models along with their associated penalties. Here, $k$ represents the clonality, and $m_{0}$ denotes the reference diploid coverage.

| **Karyotype** | **Coverage (m)** | **Allelic Imbalance (ai)** | **Penalty** |
| --- | --- | --- | --- |
| (AB)(2+n) | $\left( 1 + k \right) \cdot m_{0}$ | 0 | 2 |
| (AB)(2–n) | $\left( 1-k \right)\cdot m_{0}$ | 0 | 2 |
| A | $\frac{\left( 2-k \right)\cdot m_{0}}{2}$ | $\frac{k}{2\left( 2-k \right)}$ | 2 |
| AA | $m_{0}$ | $\frac{k}{2}$ | 3 |
| AAB | $\frac{\left( 2+k \right)\cdot m_{0}}{2}$ | $\frac{k}{2\left( 2+k \right)}$ | 2 |
| AAAB | $\left( 1+k \right)\cdot m_{0}$ | $\frac{k}{2+2k}$ | 3 |
| AAA | $\frac{\left( 2+k \right)\cdot m_{0}}{2}$ | $\frac{3k}{4+2k}$ | 4 |
| AAAA | $\left( 1+k \right)\cdot m_{0}$ | $\frac{k}{1+k}$ | 5 |
| A+AA | $\frac{\left( 1+k \right)\cdot m_{0}}{2}$ | 0.5 | 4 |
| AAB+AAAB | $\frac{\left( 3+k \right)\cdot m_{0}}{2}$ | $\frac{1+k}{6+2k}$ | 4 |
| AA+AAA | $\left( 1+\frac{k}{2} \right)\cdot m_{0}$ | 0.5 | 5 |
| AA+AAAA | $\left( 1+k \right)\cdot m_{0}$ | 0.5 | 5 |
| AA+AAB | $\frac{\left( 2+k \right)\cdot m_{0}}{2}$ | $\frac{2-k}{4+2k}$ | 4 |
| AAB+AABB | $\frac{\left( 3+k \right)\cdot m_{0}}{2}$ | $\frac{1-k}{6+2k}$ | 4 |
| AAA+AAAA | $\frac{\left( 3+k \right)\cdot m_{0}}{2}$ | 0.5 | 6 |

***Single-cell WGS analysis***

We implemented a multi-step pipeline to process paired-end sequencing data and derive allele-specific copy number profiles across single cells. The workflow begins with preprocessing of raw FASTQ files, where read pairs are concatenated and aligned to the human reference genome using Bowtie2. The resulting alignments are converted to sorted and indexed BAM files via Samtools. From these, targeted pileup files are generated at predefined genomic positions, and variant calling is performed using VarScan to produce VCF files. Each VCF is then parsed to extract single-nucleotide variants (SNVs) along with their reference and alternate allele read counts, producing allele count (.ac) files for each sample.

These allele count files are further processed to compute B-allele frequency (BAF) and total coverage per site. High-confidence SNPs are selected by intersecting with a curated list of reliable germline variants and known heterozygous sites. The BAF is calculated as the proportion of alternate reads, and variants are aggregated into fixed 5 Mb windows to compute average BAF and coverage per window. Diploid regions are used to normalize coverage, enabling estimation of relative copy number. Finally, genome-wide heatmaps of absolute BAF deviation (|BAF − 0.5|) and normalized copy number are generated across cells, with chromosomes along the x-axis and cells along the y-axis. These visualizations highlight allelic imbalance and copy number alterations, revealing clonal architecture and chromosomal abnormalities within the sample.

***Single-cell RNA-seq analysis***

Sequenced scRNA-seq samples were processed using Cell Ranger v7.1.0 (10x Genomics) with the GRCh38 v2020-A reference genome (refdata-gex-GRCh38-2020-A). The filtered gene-by-cell UMI count matrix was analyzed using Seurat v5.1.0 (2). Quality control removed cells with fewer than 200 genes or 1,000 UMIs (possible debris), more than 10,000 genes or 80,000 UMIs (possible multiplets), over 10% mitochondrial gene expression (indicative of stress or apoptosis), or identified as doublets by Scrublet v0.2.3 (3). After filtering, 6,593 cells remained. Normalization was performed using Seurat’s SCTransform function, regressing out sequencing depth and mitochondrial gene percentage. Dimensionality reduction was conducted using PCA with the top 30 principal components, and cells were clustered using a k-nearest neighbor graph and the Louvain algorithm at a resolution of 0.5. Doublets were identified with Scrublet using an expected doublet rate of 0.15, calculating doublet scores across 10 iterations, and determining the final cut-off via KMeans clustering.

To detect allele-specific CNAs based on scRNA-seq data, we used the “numbat” R-package v1.4.2 (4). For generating SNP pileup data, cellsnp-lite v1.2.3 (5) was used, and eagle v2.4.1 (6) was used for SNPs phasing. Common SNP VCF and phasing reference panels were downloaded for the 1000 Genome reference (hg38 build) from the links provided in the Numbat documentation (https://kharchenkolab.github.io/numbat/articles/numbat.html). Finally, we ran Numbat with the default parameters using the filtered gene-by-cell UMI count matrix from the scRNA-seq RDS object generated by Seurat (see above Methods), Numbat’s reference expression profile ref_hca, and allele counts data frame generated during pre-processing steps described above.

***Structure variation analysis***

CREST (7) structural variant predictions near S2K segment breakpoints were examined to identify candidate variants that matched the observed CNAs. As preliminary analysis indicated strong agreement between S2K’s segments and previously published structural variants, we restricted our initial search to predicted SVs within 1 Mb of the S2K breakpoints, and only broadened our search region if no appropriate candidates could be identified. Manual verification of candidate structural variants was conducted in IGV by examining the short-read alignments used to generate the SV calls. Counts of reads supporting each structural variant were tallied with fuzzion2 (<https://github.com/stjude/fuzzion2>).

**Results**

***NB5 results***

The NB5 neuroblastoma cell line exhibits extensive chromosomal instability, with strong concordance between S2K and cytogenetic analyses. S2K reveals a large segmental loss on chromosome 3 and a small gain at 3q, both with an estimated clonality of 0.5, indicating the presence of multiple subclones. This is supported by SKY analysis, which identifies two distinct subclones: one harboring a 1-copy loss of the entire chromosome 3 and another retaining an intact chromosome 3. Both clones show 3q gain resulting from a t(3;14)(q26.32;q22.1) translocation, consistent with the focal 3q gain detected by S2K. Additional S2K-detected abnormalities, including widespread single-copy losses on chromosomes 6, 13, and 18, and focal high-level amplifications, match the cytogenetic findings of del(6)(q22.3q26), monosomies of chromosomes 13 and 18, and the presence of double minutes. The subclonal loss of one copy of chromosome 3 identified by S2K also aligns with FISH results showing that 32% of cells carry only a single RBSN probe signal at 3p25. Collectively, these findings confirm that S2K accurately reconstructs the key clonal and subclonal chromosomal alterations identified by cytogenetic profiling in NB5, including segmental imbalances, translocations, and extrachromosomal amplifications.
