## Supplementary Figure Legend for "Seq2Karyotype (S2K): A Method for *in-silico* Karyotyping Using Single-Sample Whole-Genome Sequencing Data"

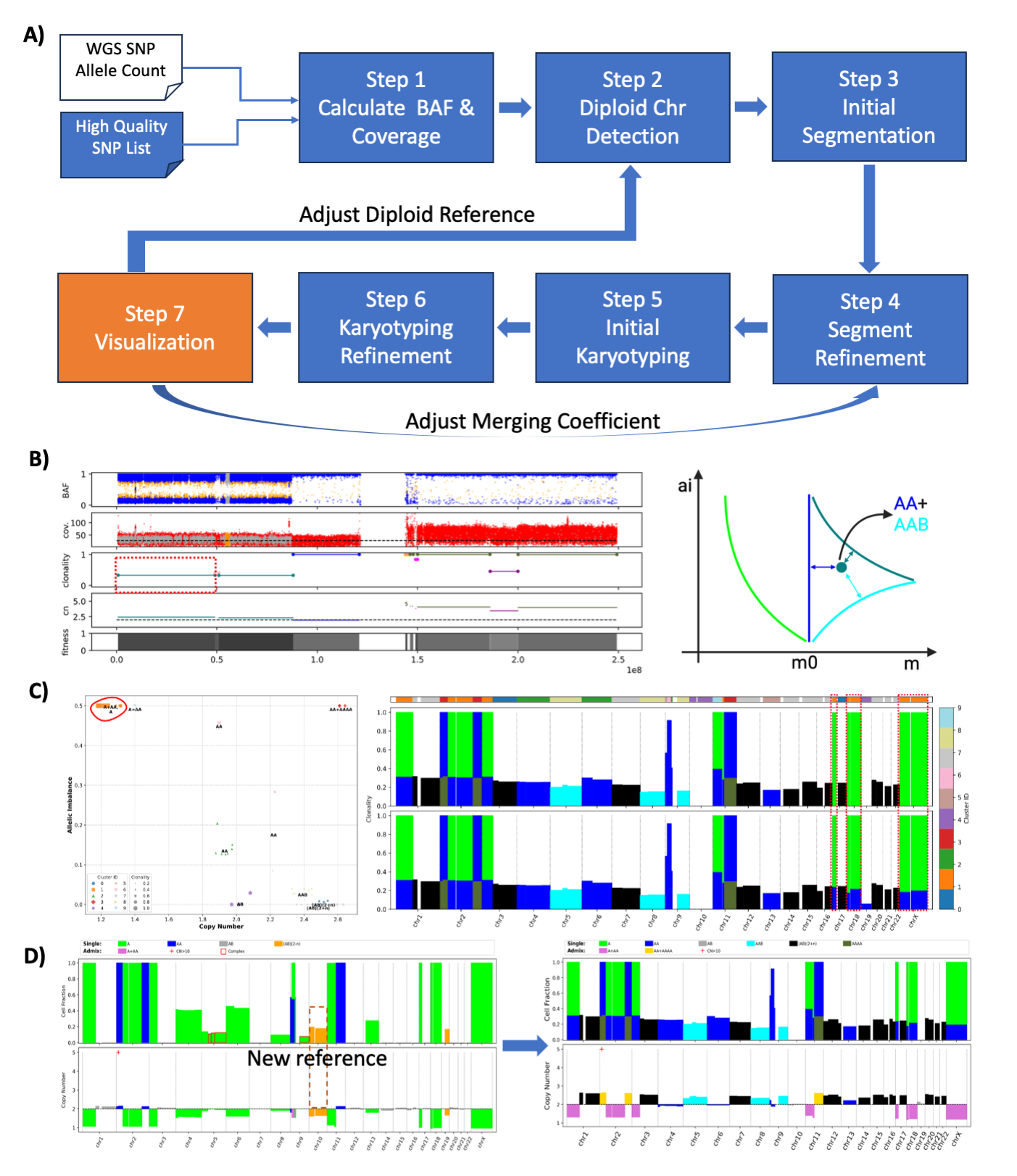

**Supplementary Figure S1: Overview of the S2K workflow and key analytical modules. (A)** Analytical process of the S2K pipeline. It starts by using allelic counts of variants derived from a single WGS sample that intersect with a high-quality SNP list to calculate B-allele fraction (BAF) and coverage. This is followed by diploid chromosome detection (Step 2), initial segmentation (Step 3), segment refinement (Step 4), initial karyotyping (Step 5) and karyotyping refinement (Step 6). The results can be visualized in the S2K karyotype viewer (Step 7) for final adjustments, allowing iterative optimization of the diploid reference and merging coefficients. **(B)** BAF, coverage, clonality, CNV segments and karyotype fitness score are visualized in left panel. Karyotyping principle is depicted in the right panel. Models for basic karyotypes: 1-copy loss (A), cnLOH (AA), 1-copy gain (AAB) are shown with lines: light green, blue and cyan respectively. Highlighted region /red rectangle/ in left panel is classified as AA+AAB because is closest to a complex model /dark green/ composed of a mixture of two karyotypes AA and AAB. **(C)** An example of clustering-based refinement of karyotype models (left) and the resulting karyotype composition across chromosomes (right). Genomic segments are grouped using Gaussian mixture clustering based on copy number and allelic imbalance, and the most likely karyotype model is assigned to each cluster by minimizing the distance to observed data. The three chromosomes with dotted red boxes (i.e. chr16, chr18, and chrX) were assigned to the karyotype model of admixture of 1-copy (loss) and 2-copy (cn-LOH) given the dominant karyotype in their corresponding clusters. **(D)** An example of reselecting a new diploid reference, showing the impact on CNV and karyotype profiles before (left, which contains the unlikely scenario of bi-allelic loss) and after (right, no bi-allelic loss) adjustment.

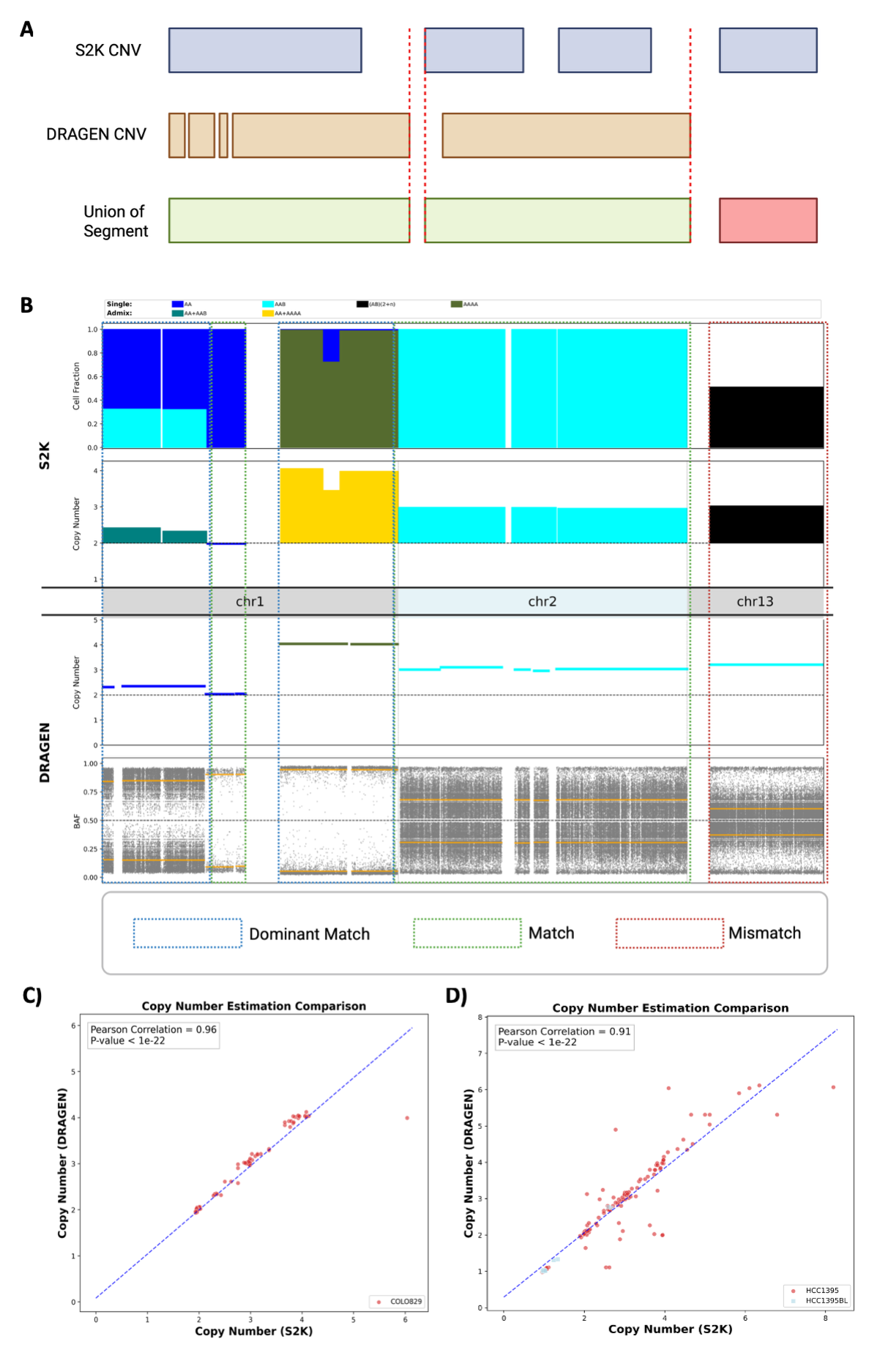

**Supplementary Figure S2: Comparison of CNV concordance between S2K and DRAGEN**. **(A)** Illustration of CNV comparison which used the union of segments computed by the two tools. Each segment is assigned with a match or mismatch status based on the consistency of the predominant karyotype (i.e. major CNV type in a region with admixed CNVs) projected by S2K versus the CNV state assigned by DRAGEN. **(B)** Illustration of CNV match status between DRAGEN and S2K in three chromosomes of COLO829. As DRAGEN does not calculate admixture of multiple CNV states, a match to the dominant CNV state predicted by S2K is considered a dominant match (dotted blue boxes) in addition to exact match (shown as dotted green boxes). Mismatches include both missed copy number events and inconsistent copy number states that are shown as the dotted red boxes. Comparison of non-diploid copy number events generated by S2K and DRAGEN is shown in **(C**) for COLO829 and in **(D)** for HCC1395 and its matched normal HCC1395BL. Only segments that are not diploid in any of the methods are shown (including cn-LOH). The matched normal of COLO829 is not shown as only diploid CNVs were detected by S2K and DRAGEN.

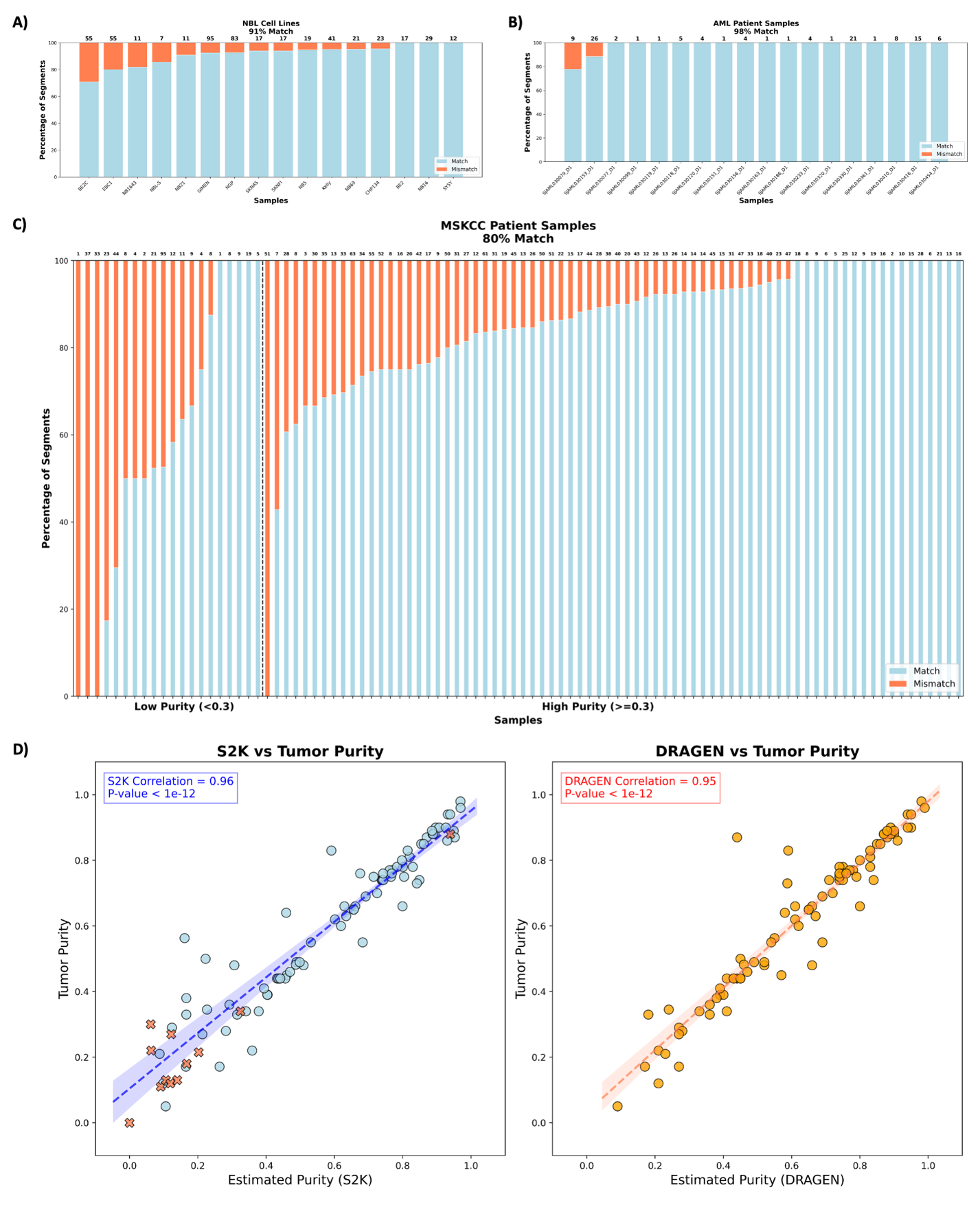

**Supplementary Figure S3: Comparison of CNV segment and patient tumor purity estimation computed by S2K vs DRAGEN.** The percentage of matched (blue) and mismatched (red) CNV segments between S2K and DRAGEN for each sample is summarized for NBL cell lines (**A**), AML samples (**B**), and (**C**) MSKCC neuroblastoma patient samples which are separated into low- and high-purity samples using a cutoff of 0.3 purity based on clinical practise (1). The number of CNV segments in each sample is displayed at the top (**D**) Comparison of published tumor purity with that estimated by S2K (left) and by DRAGEN (right). Samples marked by “X” at the left panel refer to those that only had purity estimates by S2K as DRAGEN failed to generate purity estimates for these samples.

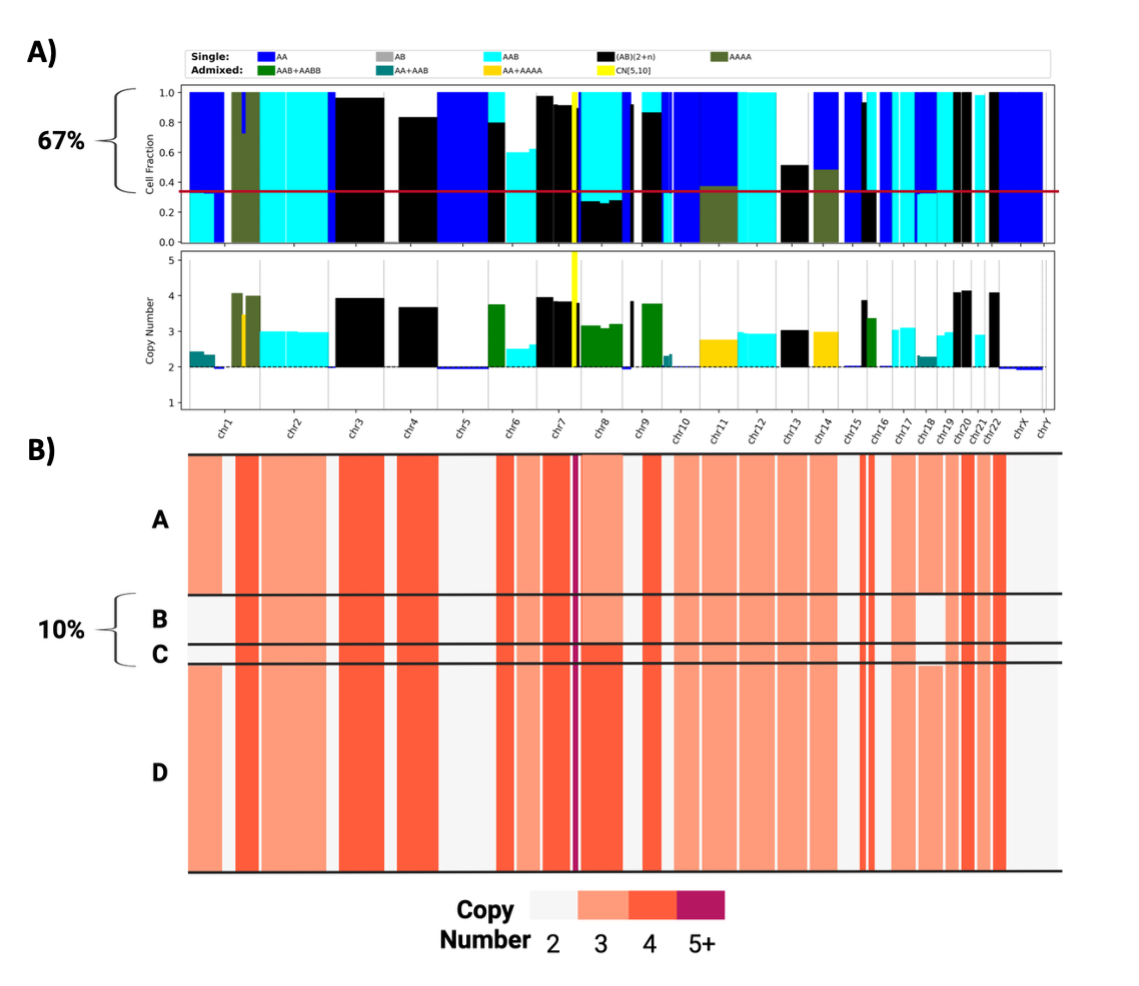

**Supplementary Figure S4: Comparison of COLO829 karyotypes based on S2K analysis with those by scWGS profiling**. **(A)** S2K karyotype view of COLO829 copy number with cellular fractions shown at the top and copy number estimates at the bottom. The red horizontal line demarcates the cellular fraction at 1p, 10p and 18 with cn-LOH. (**B**) Single-cell CNV profile of COLO829 adapted from Velazquez-Villarreal et al (2) representing the four major subgroups. The cells in subgroups B and C correspond to cn-LOH events identified by S2K at 1p, 10p and 18 while the subgroups A and D correspond to the 3-copy events representing der18 involving the three regions.

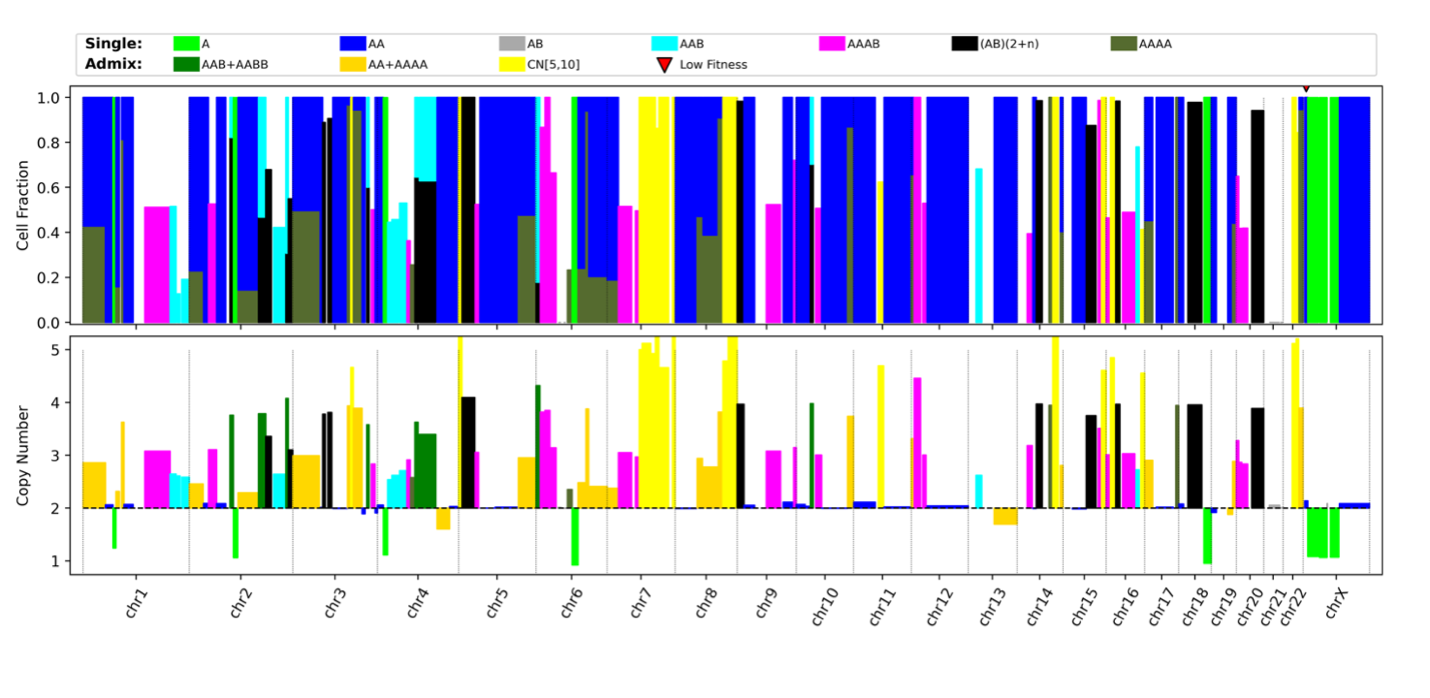

**Supplementary Figure S5: HCC1395 karyotypes based on S2K analysis.** The extensive LOH and high aneuploidy across the genome is consistent with the scWGS-based CNV profile identified by Fang et al (3).

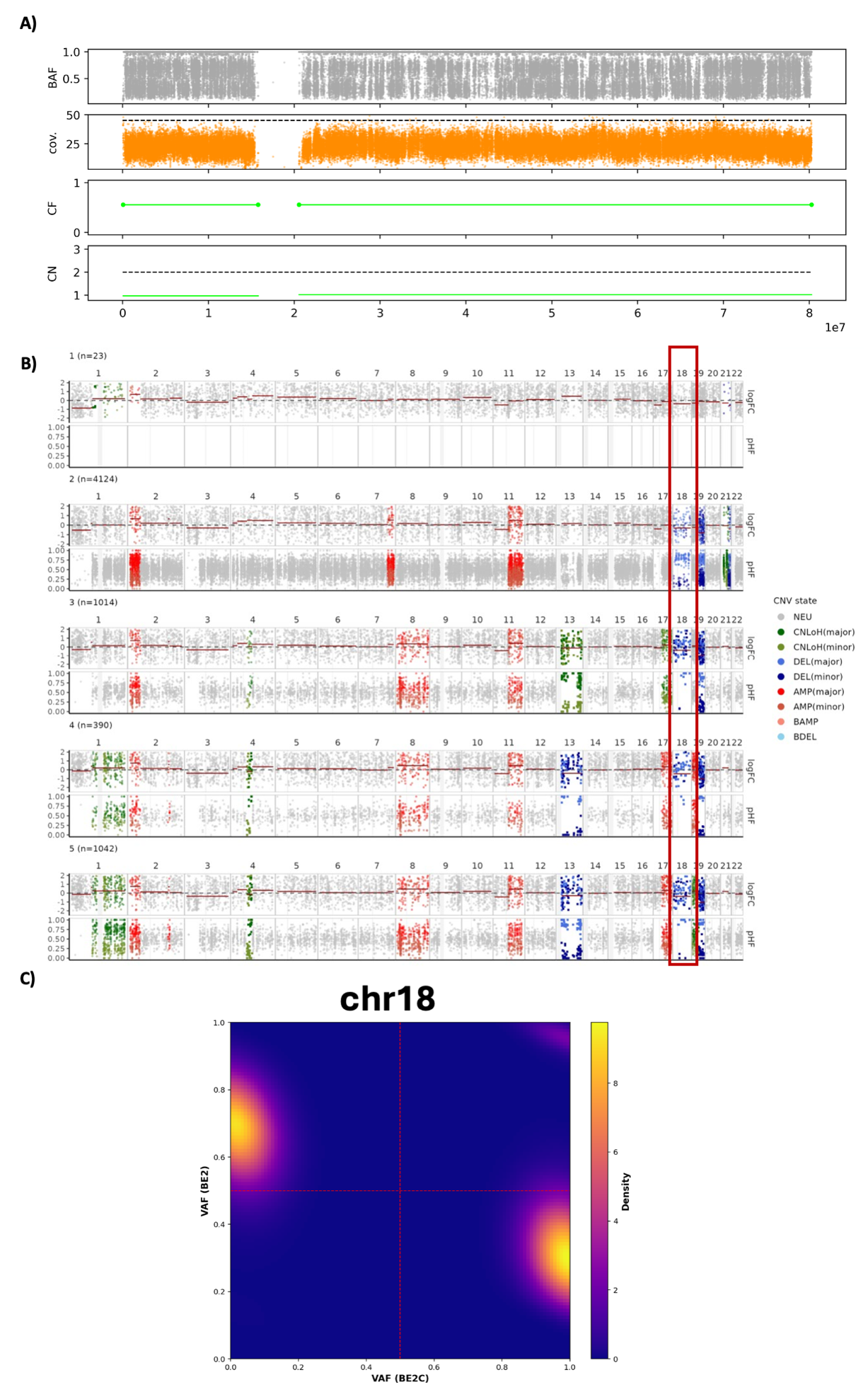

**Supplementary Figure S6: Convergent evolution of 1-copy loss of chr18 in BE2 cell line. (A)** CNV and BAF profiles of chr18 showing 1-copy CNV based on coverage analysis but incomplete allelic imbalance leading to an estimated cellular fraction of 0.6 based on BAF. **(B)** CNV profile inferred from scRNA-seq based on Numbat (4) analysis with chr18 highlighted by a red box. The largest group (group 2) which contains 4,124 single cells shows loss of different haplotypes. **(C)** Density plot comparing BAFs of chr18 SNPs in BE2C (x-axis) with those of BE2.

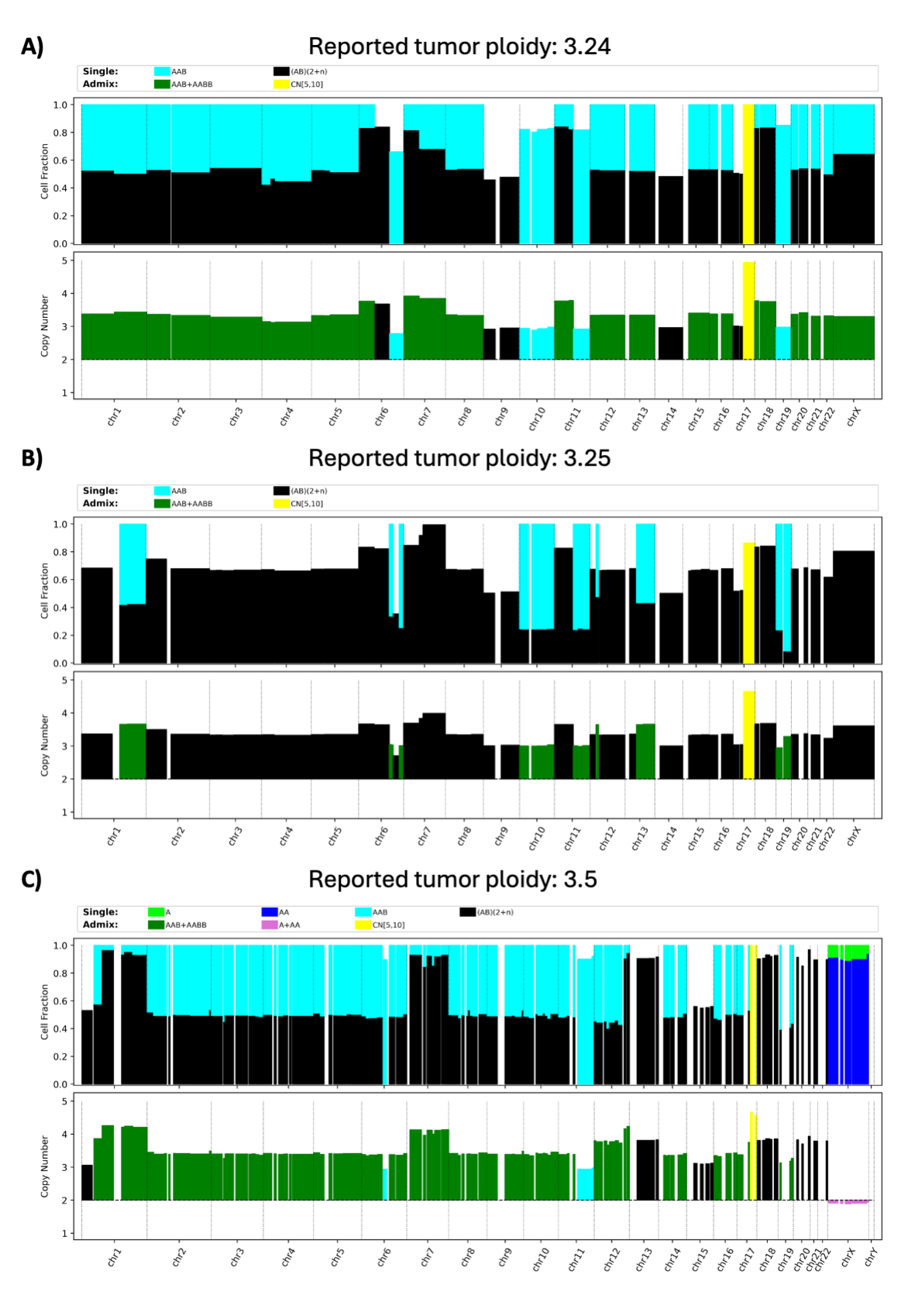

**Supplementary Figure S7: Admixture of tetraploid cells in neuroblastoma patient samples.** S2K karyotype view of three patient samples predicted to contain a subpopulation of tetraploid cells: (**A**) H-132383_D2, (**B**) H-132383_D1, and (**C**) H-136648_R1 matching the published (5) ploidies of 3.24, 3.25, and 3.5, respectively.

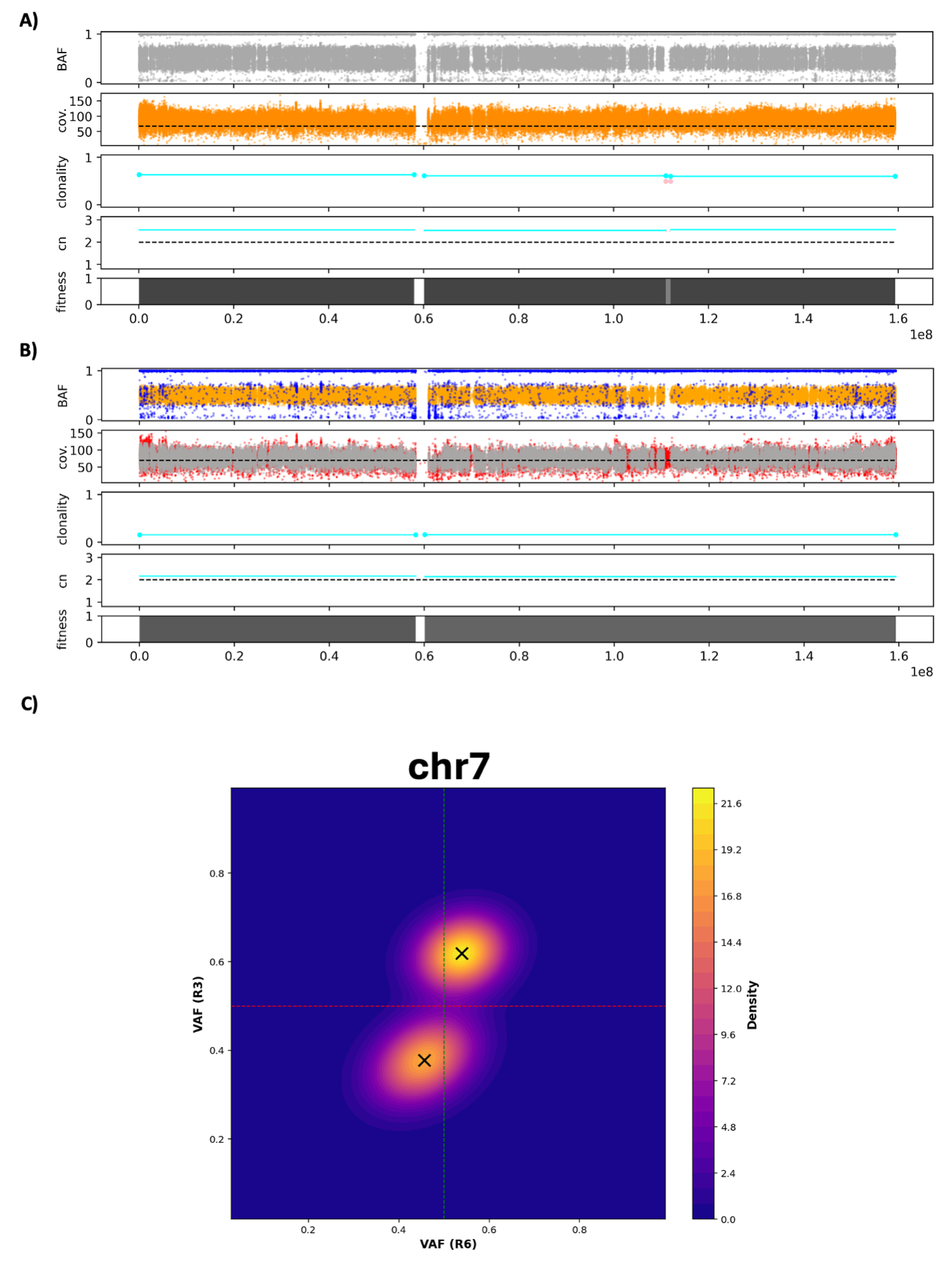

**Supplementary Figure S8: B-allele fraction (BAF) heatmap of chromosome 7 for two tumor samples derived from case H134722. (A)** S2K chromosome view of R3 and **(B)** S2K chromosome view of R6. **(C)** VAF heatmap of chromosome 7 of R3 (y-axis) vs R6 (x-axis). Both R3 and R6 contain a 1-copy gain of chr7 with predicted CF of 0.62 and 0.15, respectively. The density of BAF deviated from 0.5 is expected for a 1-copy gain and the consistency in VAF change direction in the two samples supports the amplification of the same haplotype.

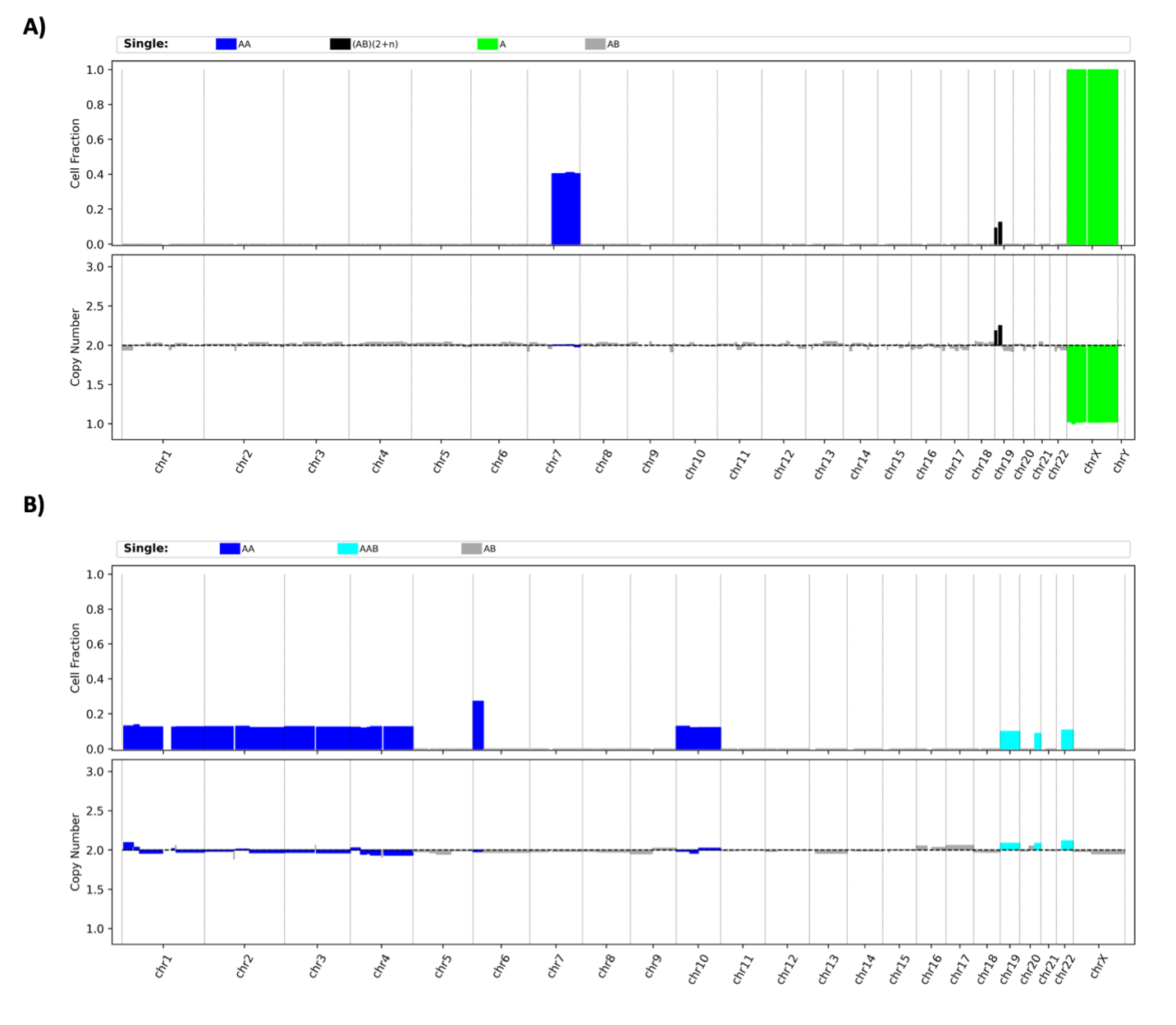

**Supplementary Figure S9: S2K analysis of two MDS cases.** S2K identified mosaic 6p uniparental disomy (UPD) and monosomy 7 in two myeloid neoplasm cases; monosomy 7 was validated by scWGS.

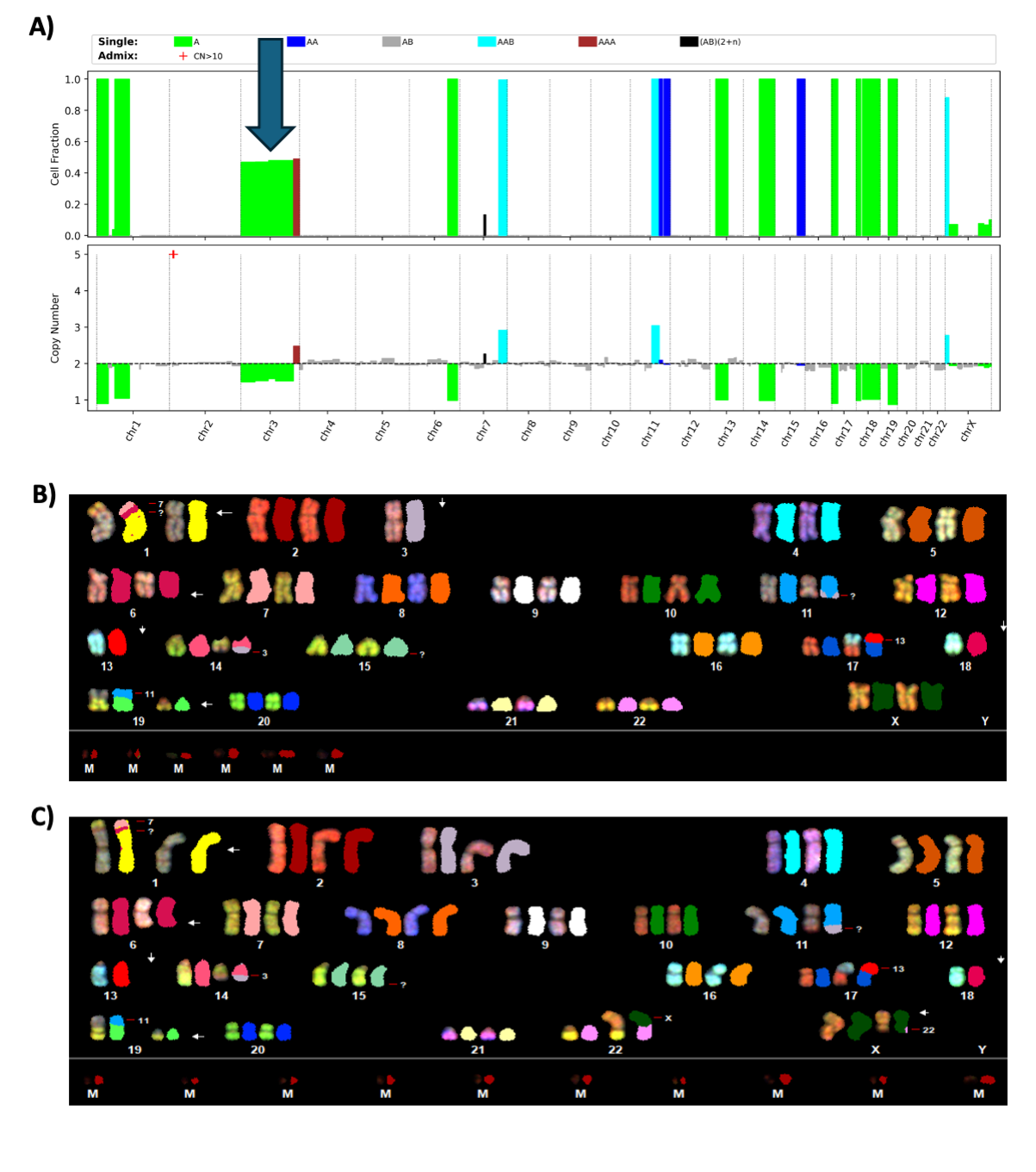
**Supplementary Figure S10: Admixture of copy-gain and copy-loss at chromosome 3 in cell line NB5. (A)** Karyotype view of S2K showing chr3 contains a large segment of 1-copy loss and a small segment at 3q of 1-copy gain with an estimated clonality of 0.5. SKY mapping validated the presence of two predicted clones in (**B**) and (**C**), both of which contain 3q gain caused by translocation between 14q and 3q. The clone shown in (**B**) has a 1-copy loss of the entire chromosome 3 while the one in (**C)** has the intact chromosome 3.
